# Escalating High-dimensional Imaging using Combinatorial Channel Multiplexing and Deep Learning

**DOI:** 10.1101/2023.09.09.556962

**Authors:** Raz Ben-Uri, Lior Ben Shabat, Dana Shainshein, Omer Bar-Tal, Yuval Bussi, Noa Maimon, Tal Keidar Haran, Idan Milo, Inna Goliand, Yoseph Addadi, Tomer-Meir Salame, Alexander Rochwarger, Christian M. Schürch, Shai Bagon, Ofer Elhanani, Leeat Keren

**Author notes:** These authors contributed equally (Raz Ben-Uri, Lior Ben Shabat).

## Abstract

Understanding tissue structure and function requires tools that quantify the expression of multiple proteins at single-cell resolution while preserving spatial information. Current imaging technologies use a separate channel for each individual protein, inherently limiting their throughput and scalability. Here, we present CombPlex (COMBinatorial multiPLEXing), a combinatorial staining platform coupled with an algorithmic framework to exponentially increase the number of proteins that can be measured from *C* up to 2^*c*^ − 1. In CombPlex, every protein can be imaged in several channels, and every channel contains agglomerated images of several proteins. These combinatorically-compressed images are then decompressed to individual protein-images using deep learning. We achieve accurate reconstruction when compressing the stains of twenty-two proteins to five imaging channels and demonstrate that the approach works in both fluorescence microscopy and in mass-based imaging. Combinatorial staining coupled with deep-learning decompression can escalate the number of proteins measured using any imaging modality, without the need for specialized instrumentation. Coupling CombPlex with instruments for high-dimensional imaging could pave the way to image hundreds of proteins at single-cell resolution in intact tissue sections.

## Introduction

Immunohistochemistry, whereby antibodies are used to quantify the single-cell spatial distribution of proteins in tissue sections, is a pillar of pathology, informing patient diagnosis, clinical decision making as well as discovery studies ^1^. Traditional chromogenic immunohistochemistry is limited to visualizing one or two proteins at a time, whereas standard immunofluorescence methods are limited to three to five proteins due to overlapping emission spectra ^2^. However, to uncover complex cellular structures, interactions, and functions, there is a need to simultaneously visualize many proteins *in situ*. To address this gap and increase the number of proteins measured in a single tissue, various multiplexed imaging approaches have been recently developed, gaining interest from both the scientific and medical communities (^3–16^ reviewed in ^17–20^).

While there are many multiplexed imaging methods, differing in the antibody tagging and detection strategies used, they can be broadly grouped into two classes. Mass-based approaches use antibodies conjugated to heavy metals, which can be resolved using a mass spectrometer ^16,21^. These methods require expensive instrumentation, and their multiplexing capability is limited to a few dozen proteins by the number of available metals for conjugation and detection. Cyclic fluorescence-based approaches increase the number of proteins measured by performing consecutive imaging cycles, each illuminating two or three proteins, followed by removal of the signal by chemical or enzymatic reactions ^3–14^. These methods are also effectively limited since the number of imaging cycles increases linearly with the number of proteins visualized, which in turn can result in tissue deterioration and prohibitively long imaging times. To date, there remains a need to increase the multiplexing capacity in imaging for any instrument.

One common characteristic of existing multiplexed imaging approaches is the use of a separate channel (e.g., fluorescent color, mass tag etc.) for each individual protein, inherently limiting the scalability of these methods (**Fig. 1A**). Combinatorial approaches, whereby each target is encoded by several channels (e.g., a sequence of colors over multiple rounds), can dramatically increase throughput (**Fig. 1B**). Such methods effectively generate a target-specific barcode and have been successfully used to measure thousands of mRNAs ^22,23^. One assumption that lies at the basis of these combinatorial approaches is the ability to optically resolve individual molecules (‘spot counting’) to identify the underlying barcodes. This assumption can be upheld for the mRNAs of many genes if using a high enough magnification (400-1000X). If this assumption is invalidated and barcodes spatially overlap, it becomes challenging to differentiate between different genes ^23^. Since proteins are on average 10,000-fold more abundant than mRNAs ^24^, the spot counting assumption is invalidated for most proteins under standard magnifications, and therefore to date combinatorial staining methods have not been applied to imaging proteins.

**Figure 1:**
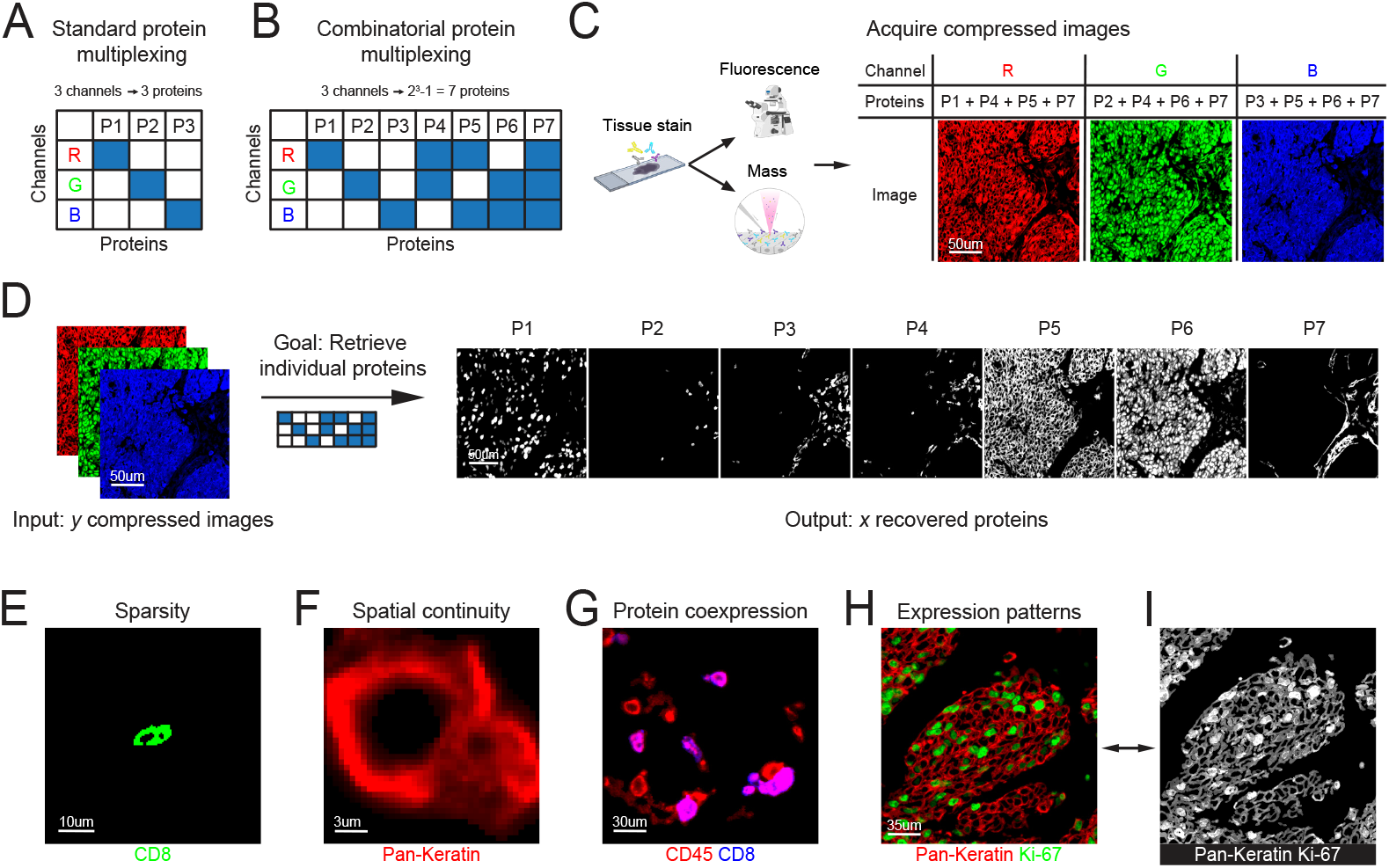
Combinatorial channel multiplexing escalates protein imaging. **(A)** In standard protein imaging, the image of each protein (columns) is acquired on a separate measurement channel (e.g. fluorescent color, rows). **(B)** In combinatorial protein multiplexing, the image of each protein (columns) is acquired on a unique combination of measurement channels (rows). This increases the number of proteins that can be measured using a given number of channels exponentially. **(C)** Illustration of the acquisition of combinatorically-compressed images using fluorescent or mass-based approaches. Each channel in the image depicts the signal of multiple proteins. **(D)** Given compressed images acquired as detailed in (B-C) and the combinatorial staining matrix, the objective is to reconstruct the underlying single-protein measurements. **(E-I)** Examples of properties of protein-images that can serve as constraints during signal reconstruction. **(E)** CD8 exhibits sparse staining in tissues. **(F)** Keratins exhibit continuous signal. **(G)** CD8 (blue) is co-expressed with CD45 (red). **(H)** Pan-Keratin (red) exhibits cytoplasmic staining while Ki-67 exhibits nuclear staining (green). **(I)** Same as in (H). Although both proteins are shown in grayscale, they can be differentiated by their unique staining patterns.

Mathematically, resolving the spatial distribution of *P* proteins from combinatorial staining using *C* channels (*P* ≫ *C*) while allowing spatial overlap is underdetermined, meaning that it has an infinite number of possible solutions (see details in the results). Work in signal processing has shown that solutions for underdetermined linear systems can be recovered if the underlying data is structured. For example, in compressed sensing ^25^, the assumption that the underlying data is sparse is often used to constrain the infinite plausible solution space. Previous work has leveraged the structure of gene expression modules to apply compressed sensing to increase the throughput of both single-cell RNA sequencing ^26^ and single molecule mRNA imaging ^27^. However, one limitation of compressed sensing is that it explicitly formulates the constraints on the solution space, and as such is highly sensitive to the predefined structure of the data. Thus, these methods require elaborate setups such as performing single cell RNA sequencing (scRNASeq) prior to imaging, which is expensive and often incompatible with the availability of clinical material. While similar approaches have been suggested for proteins ^28^, in effect they have never materialized beyond simulations of cells in suspension, presumably due to the inherent limitations of the method.

In this work we describe CombPlex (COMBinatorial multiPLEXing), an experimental foundation for combinatorial staining of proteins in tissues coupled with an algorithmic framework for accurately decompressing the combined signal back to individual proteins. In CombPlex, each protein can be measured in several channels, and each acquired image captures a combination of signals from several proteins. CombPlex utilizes a deep neural network to learn meaningful priors on the underlying protein images and reconstruct individual protein images from combinatorically-compressed measurements. We devise experimental protocols to perform combinatorial staining and show that we can accurately multiplex multiple proteins on a single channel. We demonstrate accurate (F1>0.97, R>0.93) reconstruction when staining for seven proteins using three fluorescent channels, indicating that the approach is compatible with conventional fluorescent microscopes that are widely used in research and in the clinic. We further test it in cyclic fluorescence (CODEX) and mass-based imaging (MIBI), using up to 5 channels to accurately measure 22 proteins, showing that the approach is applicable to a wide variety of platforms and tissue specimens. Using simulations, we show that the neural network learns several degrees of structure in the data including the compression scheme, protein co-expression and spatial expression patterns. Altogether, we demonstrate how combinatorial staining coupled with deep-learning decompression can serve to escalate the number of proteins measured using any imaging modality, without the need for specialized instrumentation. Coupling CombPlex with instruments for high-dimensional imaging can pave the way to imaging hundreds of proteins at single-cell resolution in intact tissue sections.

## Results

### Combinatorial channel multiplexing escalates protein imaging

We formally define CombPlex as a method for measuring *P* proteins using *C* measurement channels, where *P* ≫ *C*. A measurement channel can refer to, e.g., a particular color in fluorescent microscopy or a particular mass range in mass-based imaging. Each protein is measured using a unique combination of channels. This results in *compressed images*, in which each channel contains a combination of the signals of several proteins (**Fig. 1B,C**).

Formally, the multiplexed-image compression can be estimated as:

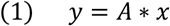

where *y* is a multiplexed-image, of dimensions height x width x channels, with *C* channels. *A* defines the compression matrix of size *C* × *P, x* is the set of *P* single-protein images and ∗ is the convolution operator (Methods). This staining scheme exponentially increases the number of proteins that can be imaged using a fixed number of channels. By exploiting all possible combinations, we get *P* = 2^*c*^ − 1. So, for example, using three-color fluorescent microscopy one can measure seven proteins (2^3^ − 1) as illustrated in figure 1B. A multiplexed imaging instrument with forty channels could potentially measure 10^12^ targets, orders of magnitude more than needed to measure the entire proteome. Although this is limited by experimental and practical considerations, such as antibody availability, cost and the quality of staining, it could drastically increase multiplexing capabilities.

Given a set of compressed images *y* and a pre-defined compression matrix *A*, we want to recover the original protein images, i.e., solve for *x* **(Fig. 1D)**. Altogether this defines a set of *C* linear equations with *P* variables for each pixel. Since *P* ≫ *C*, this problem is underdetermined and has a degenerate solution space with an infinite number of solutions.

We reasoned that although the mathematical problem of compressing protein images is underdetermined, additional properties of the images may be leveraged to constrain the solution space and recover the true *x*. For example, images of proteins are often sparse, with only a minority of pixels containing a signal for any particular protein (**Fig. 1E**). Moreover, protein expression is continuous in space. If a pixel stains positive for a particular protein, it is likely that nearby pixels will also stain positive for that same protein (**Fig. 1F**). In addition, protein expression is structured, with particular proteins tending to cooccur together or be mutually exclusive. For example, pixels that stain positive for CD8, expressed on T cells, will also stain positively for CD45, expressed on all immune cells, but not vice-versa (**Fig. 1G**). Finally, different proteins vary in their staining patterns (e.g., membranous vs. nuclear) and their overall abundance (strong vs. weak staining) (**Fig. 1H**). The unique staining pattern of proteins enables a trained person to recognize a variety of widely used antibodies, even without labels. For example, a skilled biologist can visually separate the signals of Ki-67, a nuclear protein, from keratins, which are expressed in the cytoplasm, even when both are stained together in a single image (**Fig. 1I**). Therefore, it should be equally possible to train an algorithm to do the same. We hypothesized that the various layers of structure in biological images could serve as priors to accurately reconstruct the images of single proteins from combinatorically-compressed images, even though equation (1) is underdetermined.

### *CombPlex*, a deep learning approach to decompress combinatorically-stained images, accurately reconstructs compressed images in simulations

To devise a computational framework to decompress combinatorically-acquired images, we first performed *in silico* simulations. We downloaded a published dataset, in which 57 proteins were measured in colorectal cancer (CRC) specimens using co-detection by indexing (CODEX) cyclic fluorescence ^29^. From these data, we chose 41 fields of view (FOVs) and 22 proteins that had specific staining, with good signal to noise ratio. We then simulated combinatorial measurements *in silico* by a linear combination of the protein-images according to the compression matrix illustrated in figure 2A. The matrix was designed to minimize the partition of proteins between channels and balance the number of proteins compressed in each channel (additional considerations are detailed in the methods). Overall, this resulted in 41 multiplexed images, each containing 5 channels, depicting the expression of 22 proteins (**Fig. 2B**, Methods).

**Figure 2:**
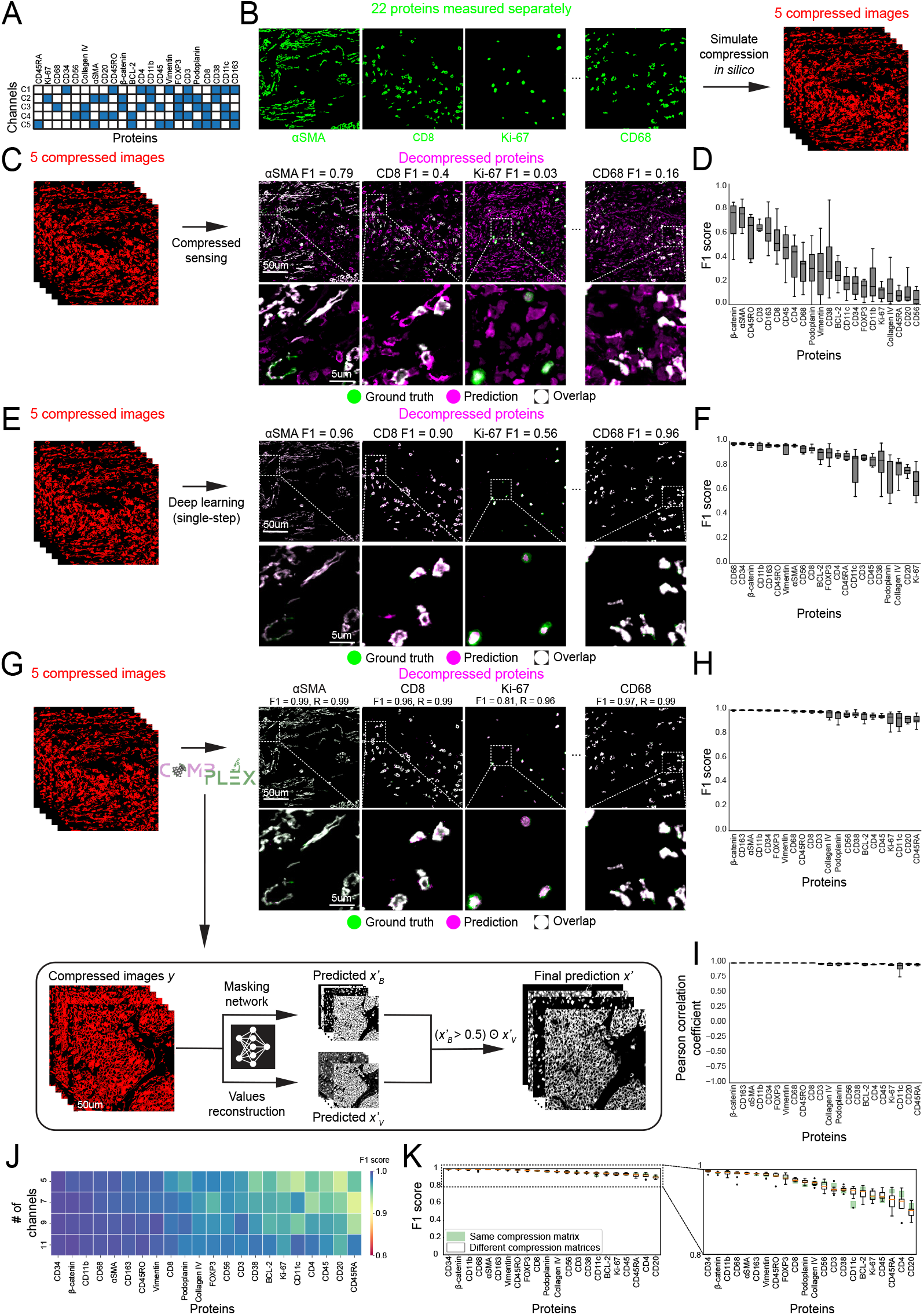
*CombPlex*, a deep learning approach to decompress combinatorically-stained images, accurately reconstructs compressed images in simulations. **(A)** The compression matrix employed to condense the data from Schürch et al ^29^. **(B)** *In silico* compression of 22 protein images into 5 combinatorically-compressed images. **(C)** A combinatorically-compressed image (red) is decompressed into individual protein images using compressed sensing. Accuracy is evaluated by calculating the F1 scores of the recovered images (magenta) compared to experimentally-measured single protein images (ground truth, green). White pixels were accurately recovered, whereas green and magenta indicate false negatives and false positives, respectively. **(D)** F1 scores (y-axis) for each one of the 22 proteins (x-axis) recovered by compressed sensing on eight test FOVs. **(E)** Same as (C), using a convolutional neural network (CNN) for reconstruction. The network uses MSE loss to recover the signal in a single step **(F)** F1 scores (y-axis) for each one of the 22 proteins (x-axis) recovered by a single-step CNN on the same eight FOVs. **(G)** Same as (C), using the CombPlex algorithmic framework for reconstruction. CombPlex is designed as a two-step process, combining a CNN that outputs binary predictions, and a CNN that recovers the values. **(H)** F1 scores (y-axis) for each one of the 22 proteins (x-axis) recovered by CombPlex on the same eight FOVs. **(I)** Pearson correlation coefficients (y-axis) between the ground truth images and the images recovered by CombPlex for the 22 proteins (x-axis) on the same eight FOVs. **(J)** Simulations were performed to compress the 22 proteins (x-axis) into 5, 7, 9 or 11 channels (y-axis) and models were trained accordingly. Colors denote the median F1 scores across eight FOVs for the individual protein images recovered using CombPlex. **(K)** Ten models were trained using compression matrix A, while shuffling the columns to change the proteins’ location in the matrix. White boxplots show the distributions of median F1 scores in the ten models across eight FOVs (y-axis) for each of the 22 proteins (x-axis). Green bars mark the 25^th^ and 75^th^ percentiles of the median F1 scores of 5 models trained with the same compression matrix. Zoom (right) shows the range between 0.8-1.

We tested different approaches to decompress the simulated images and recover the original input images. We first tried to use compressed sensing, following literature that demonstrated success for similar problems in the analysis of gene expression data (Methods ^26,27^). However, using constraints for sparsity and spatial signal continuity, compressed sensing failed to accurately recover the images of individual proteins (Median F1 scores of 0.28±0.26, R=0.51±0.33, PSNR=36.72±7, SSIM=0.92±0.08 **Fig. 2C,D, Supp. Fig. 2A**). We reasoned that more informative priors on the patterns and properties of protein expression could address the problem, but such priors are difficult to formulate mathematically. We therefore turned to deep learning, in which a neural network can leverage intricate structures of the underlying data to reconstruct the single-protein images.

We posed the problem as a supervised learning task and used deep learning to solve it. We trained a convolutional neural network (CNN), which, given a compressed image with *C* channels (*y*), recovers the underlying single-protein images, *x* (Methods). We trained the network on the simulated compressed images described above. We used 33 random FOVs for training and evaluated the results on 8 images. This approach dramatically improved the overall results (F1 = 0.91±0.16, R=0.993±0.04, PSNR=57.22±8.97, SSIM=0.9992±0.002 **Fig. 2E,F, Supp. Fig. 2A**), but performance varied significantly between proteins and FOVs. Investigation of the errors over training suggested that the network had a high incentive to maximize the reconstruction of highly abundant signals. From a biologist’s point of view, false predictions of positive and negative signals are often more crucial than the precise reconstruction of abundant signals (see elaboration on this point in the methods). We therefore implemented a hybrid approach, using the intersection of two U-Nets, one which predicts where different proteins are expressed in the image, and the other which aims to recover the actual intensity values (**Fig. 2G**). The complete pipeline is described in the methods section and illustrated in supplementary figure 1.

We tested CombPlex on the simulated compressed images described above. We used 33 random images to train the CNN and evaluated the results on 8 images. We found that CombPlex achieved excellent results, with median F1 scores of 0.98±0.11, dramatically outperforming the reconstruction using compressed sensing or deep learning alone (**Fig. 2G**,**H**). CombPlex also achieved high correlation between the reconstructed values and the values measured in the ground truth images (Median Pearson R=0.994±0.03, PSNR=57.8±8.82, SSIM=0.9993±0.002, **Fig. 2I, Supp. Fig. 2A**). We also tested CombPlex in a 4-fold cross-validation setting, each time testing on a different subset of 8-9 images and achieved highly accurate reconstruction, with median F1 scores of 0.98±0.13 across channels, and Pearson correlation coefficients of 0.994±0.1, PSNR of 59.31±10.02 and SSIM of 0.9996±0.002 across all images in the dataset (**Supp. Fig. 2B-E**). The accuracy of reconstruction was weakly related to the abundance and intensity of the signal (R= 0.24, p= 0.004, **Supp. Fig. 2F**,**G**), suggesting improved results for highly-expressed proteins that appear more frequently in the dataset. Performance was not affected by the subcellular localization of the protein, suggesting that the approach is compatible with a large variety of proteins.

Next, we evaluated the sensitivity of CombPlex to the properties of the compression. In the simulations, 22 proteins were compressed to 5 channels, which is the minimal number of channels needed to encode 22 proteins. We tested whether relaxing the level of compression by increasing the number of channels affects CombPlex’ performance. To this end, we simulated compression of the same 22 proteins to either 5, 7, 9 or 11 channels. We found that increasing the number of channels generally increased the accuracy of the reconstruction. Overall, the mean F1 for five channels was 0.95±0.11 and increased to 0.97±0.10 for 11 channels (**Fig. 2J**). We conclude that there is a tradeoff between the degree of compression and accuracy of reconstruction, and that the compression can be tailored to achieve a desired level of reconstruction accuracy.

There are many options to compress 22 proteins to 5 channels. We therefore generated 10 permutations of the compression matrix *A*, for each one simulated compressed data and then evaluated CombPlex’ performance in recovering the stains of the individual proteins. We found that CombPlex is relatively robust to the specific compression scheme used, though some matrices performed better than others (**Fig. 2K**). Standard errors for the median F1 scores of individual channels in different permutations of *A* ranged from 0.0004 to 0.008, similar to those obtained from retraining the network without changing *A*. We conclude that CombPlex can accommodate different compression matrices, allowing *A* to be largely determined by experimental considerations, such as availability of specific antibodies on specific channels etc.

Altogether, *in silico* simulations suggested that it is possible to compress the stains of many proteins to fewer measurement channels, and then recover the signal using deep learning.

### CombPlex accurately reconstructs compressed cancer images measured by fluorescent microscopy

Following successful performance of CombPlex in simulations, we proceeded to test it experimentally. Most fluorescent imaging platforms allow measuring three channels, so we decided to test a compression scheme of seven proteins to three channels, which is the maximal level of compression. We chose seven target proteins with clinical relevance that are routinely used in immunostaining of tumors: Pan-Keratin, αSMA, Ki-67, p53, CD45, CD8 and HLA-DRDPDQ (HLA-II). These proteins constitute a variety of strong and weak, nuclear, cytoplasmic and membrane, and tumor and immune stains, and are a good representation of a plausible biologically-relevant immune-oncology panel. Importantly, this panel is expected to generate images with colocalization of proteins. For example, we expect to find pixels that are positive for CD45 and CD8 (cytotoxic T cells), Ki-67 and p53 (proliferating tumor cells), CD45 and HLA-II (myeloid cells) etc. Altogether, the selection of targets for the panel, and the compression scheme, utilizing all degrees of freedom in A, represent a realistic and difficult test case for the approach (**Fig. 3A**).

**Figure 3:**
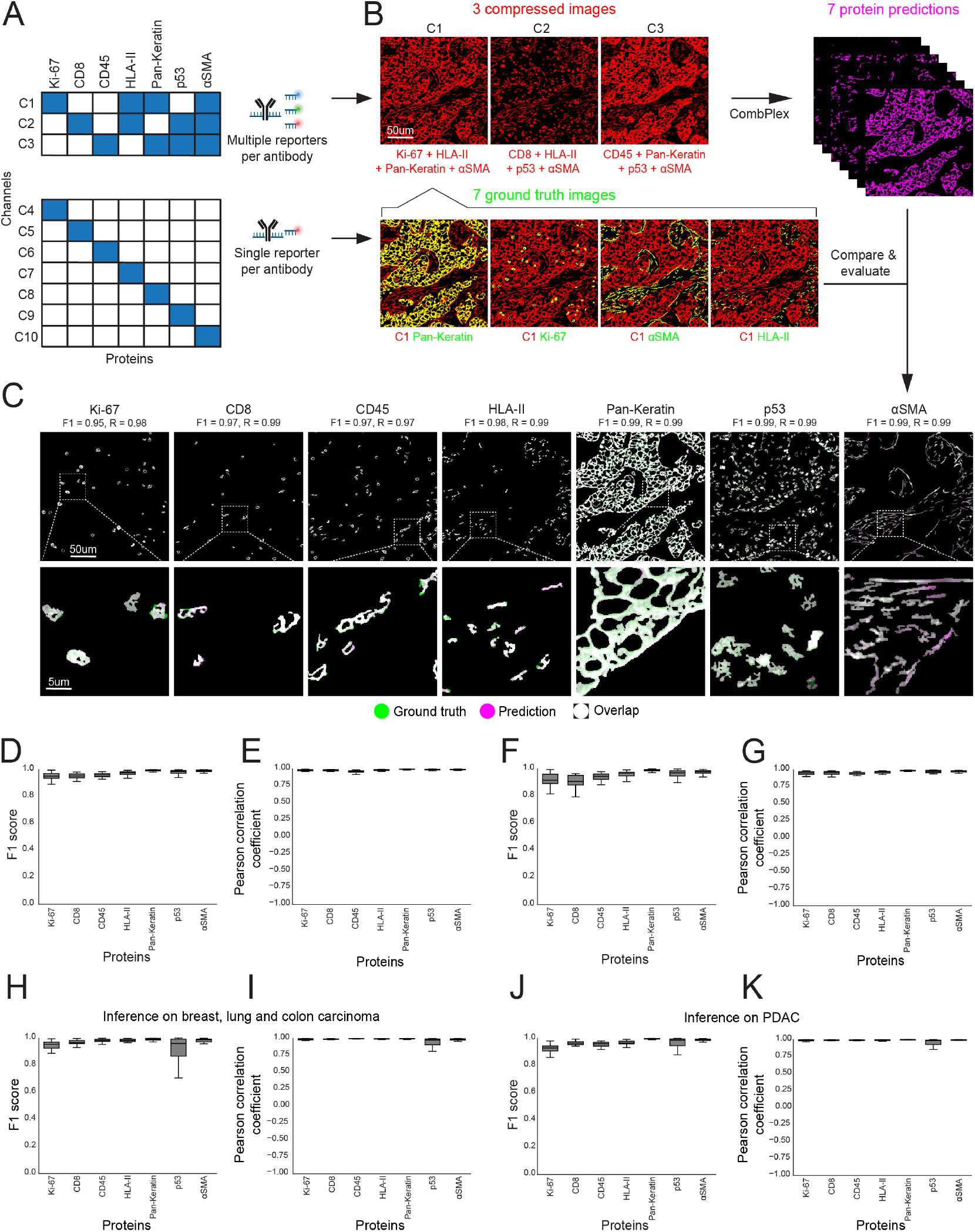
CombPlex accurately reconstructs compressed cancer images measured by fluorescent microscopy. **(A)** Experimental layout. CODEX cyclic imaging was used to image seven proteins on a single tissue section, both compressed to three measurement channels using multiple reporters per antibody (top) and individually, using a single reporter per antibody (bottom). **(B)** Top: For a single FOV, shown are the three combinatorically-compressed images (red), decompressed by CombPlex to seven reconstructed single-protein images (magenta). Bottom: overlay of four experimentally-measured single-protein images (ground truth, green) with the compressed image that contains all of their signals (red). Yellow pixels indicate overlap. **(C)** CombPlex reconstruction (magenta) is overlaid with the ground truth single-protein images (green). White pixels indicate overlap. **(D)** F1 scores (y-axis) for each protein (x-axis) recovered by CombPlex on 32 test FOVs. The models were trained on experimentally-compressed images. **(E**) Same as (D), showing the Pearson correlation coefficients between the reconstructed and ground truth images. **(F)** Same as (D). The models were trained on simulated compressed images and evaluated on experimentally-compressed images. **(G)** Same as (E). The model was trained on simulated compressed images and evaluated on experimentally-compressed images. **(H-I)** CombPlex was trained on images of breast, lung and colon carcinoma, and was tested on 44 breast, lung and colon carcinoma specimens. For each protein (x-axis), ground truth single stains were compared to CombPlex reconstructions and evaluated using F1 score (H) and Pearson correlation (I). **(J-K)** Same as (H-I) when the model trained on breast, lung and colon carcinoma was evaluated on 20 PDAC samples.

We designed a matrix *A* to compress the staining of the seven target proteins (P = 7), into three images (C = 3). Experimentally, this is achieved by labeling several antibodies with the same fluorophore. We then used CO-Detection by IndEXing (CODEX) to measure on a single tissue section both the ground truth images of individual targets and the compressed images, where the same targets are measured in a combinatorial manner (**Fig. 3A**, Table S1, Methods). Altogether the experiment required ten channels: three for the compressed images and seven for the individual targets. We stained and imaged 132 breast cancer samples, from 66 different patients (**Supp. Fig. 3A**). We performed several analyses to gauge our protocol. First, we validated that different antibodies could be measured using the same exposure, and that this did not cause a loss in signal of any respective antibody (**Supp. Fig. 3B**,**C**). Next, we evaluated that the compressed images indeed contained signals overlapping with the expected proteins and did not contain signals of other proteins in the panel. For example, the ATTO-550 channel contained signal matching the single-channel images of Keratin, Ki-67, αSMA and HLA-II, but did not contain signal matching p53, CD8 or CD45 (**Fig. 3B, Supp. Fig. 3D**). Moreover, pixel values in the compressed images were well-correlated with a simulation of a compressed channel, generated by overlaying the corresponding images in which the proteins were measured individually (Methods, **Supp. Fig. 3E**,**F**). Altogether, we concluded that it is experimentally possible to multiplex the stains of several antibodies on a single measurement channel.

We evaluated CombPlex’s performance in recovering the signal of the seven individual proteins from the compressed images. We trained the network on pairs of compressed images (*y*) and their corresponding individual-protein images (*x*). We used 100 paired images for training and evaluated performance on 32 held-out images. We found that CombPlex achieved highly accurate reconstruction, with median F1 scores on the test data of 0.98±0.07 across channels, and Pearson correlation coefficients of 0.99±0.02, PSNR of 53±9 and SSIM of 0.997±0.009 (**Fig. 3C-E, Supp. Fig. 3G**,**H**). Calculating Pearson’s correlation only on the intersecting pixels of both images was also very high (R = 0.97±0.05, **Supp. Fig. 3I**). We also tested CombPlex in a 4-fold cross-validation setting, each time testing on a different subset of 32-35 images and achieved highly accurate reconstruction, with median F1 scores of 0.98±0.09 across channels, and Pearson correlation coefficients of 0.99±0.05, PSNR of 53±10 and SSIM of 0.998±0.007 across all images in the dataset (**Supp. Fig. 3J-M**).

Next, we wondered whether we could circumvent the need to obtain training data in the form of compressed measurements acquired on the same tissue as the ground-truth single-protein measurements. Removing this requirement would be beneficial in making the approach more accessible, since pairs of compressed and single-protein images are not widely available and would have to be generated for each set of proteins and the specific compression matrix A. Simulating compressed images from widely-available sets of single-protein measurements would negate the need to acquire such data, and would also allow flexibility by enabling to use the same training data for different compression schemes.

We therefore devised a scheme to simulate compressed images (*y*) from measurements of individual proteins (*x*) according to equation (1). For these simulations, we performed augmentations that are relevant to the domain measurement system. For example, we added noise to the compression matrix (*A*) to account for experimental differences in intensities between fluorescent channels. Additional augmentations are detailed in the methods section *Simulation of compressed images*. We then trained CombPlex again, this time on pairs of individual protein images (*x*) and simulated compressed images (*y* = *sim*(*x*)). We then evaluated CombPlex’s performance on real, experimentally-compressed images. We found that even though it was trained on simulations, CombPlex achieved a highly accurate reconstruction, with median F1 scores on the test data of 0.96±0.1 across channels, Pearson correlation coefficients of 0.97±0.06, PSNR of 49±12 and SSIM of 0.993±0.014, largely comparable to the results achieved when training on real compressed images (**Fig. 3F-G, Supp. Fig. 3N**,**O**). We conclude that it is possible to train CombPlex on simulations of compressed images generated from images in which the targets were measured individually, which is valuable for the utility and applicability of the approach.

Finally, we tested the ability of CombPlex to generalize over different tissues. To this end, we trained one model on 166 samples of breast cancer, colorectal cancer and lung cancer **(Fig. 3H)**. We tested the performance of CombPlex on a heldout test set of 44 compressed images of the same tumor types. We found that CombPlex performed very well on this dataset, indicating that it is possible to train a single model for several tissue types (F1=0.98±0.11, R=0.992±0.094, PSNR=57±10, SSIM=0.998±0.006, **Fig. 3H,I, Supp. Fig. 3P**,**Q**). We then tested the model on 20 pancreatic ductal adenocarcinoma (PDAC) samples, which the network did not train on. We found that CombPlex performed very well on this novel tissue type, equivalent to its performance on heldout tissues of breast, colorectal and lung cancer (F1=0.98±0.04, R=0.992±0.05, PSNR=55±9, SSIM=0.998±0.004, **Fig. 3J**,**K, Supp. Fig. 3R**,**S**). We conclude that CombPlex can generalize across tissues for proteins with similar staining patterns.

### CombPlex can support large-scale compressions of 10 antibodies per channel

Following successful establishment of a small-scale compression scheme of seven antibodies in three channels, we evaluated the performance of the approach in a larger-scale compression, which necessitates to multiplex more proteins in each fluorescent channel. We chose 22 target proteins and designed an experiment to measure them in five channels. This scheme multiplexes 9-10 proteins per channel and represents a 4.5-fold compression (**Fig. 4A**). We utilized the CODEX system to obtain both the compressed and ground-truth images on the same breast cancer TMA. We validated that the compressed images indeed contained signals overlapping with the expected proteins and did not contain signals of other proteins in the panel (**Fig. 4B, Supp. Fig. 4A**). Altogether, we concluded that this larger-scale compression is experimentally feasible.

**Figure 4:**
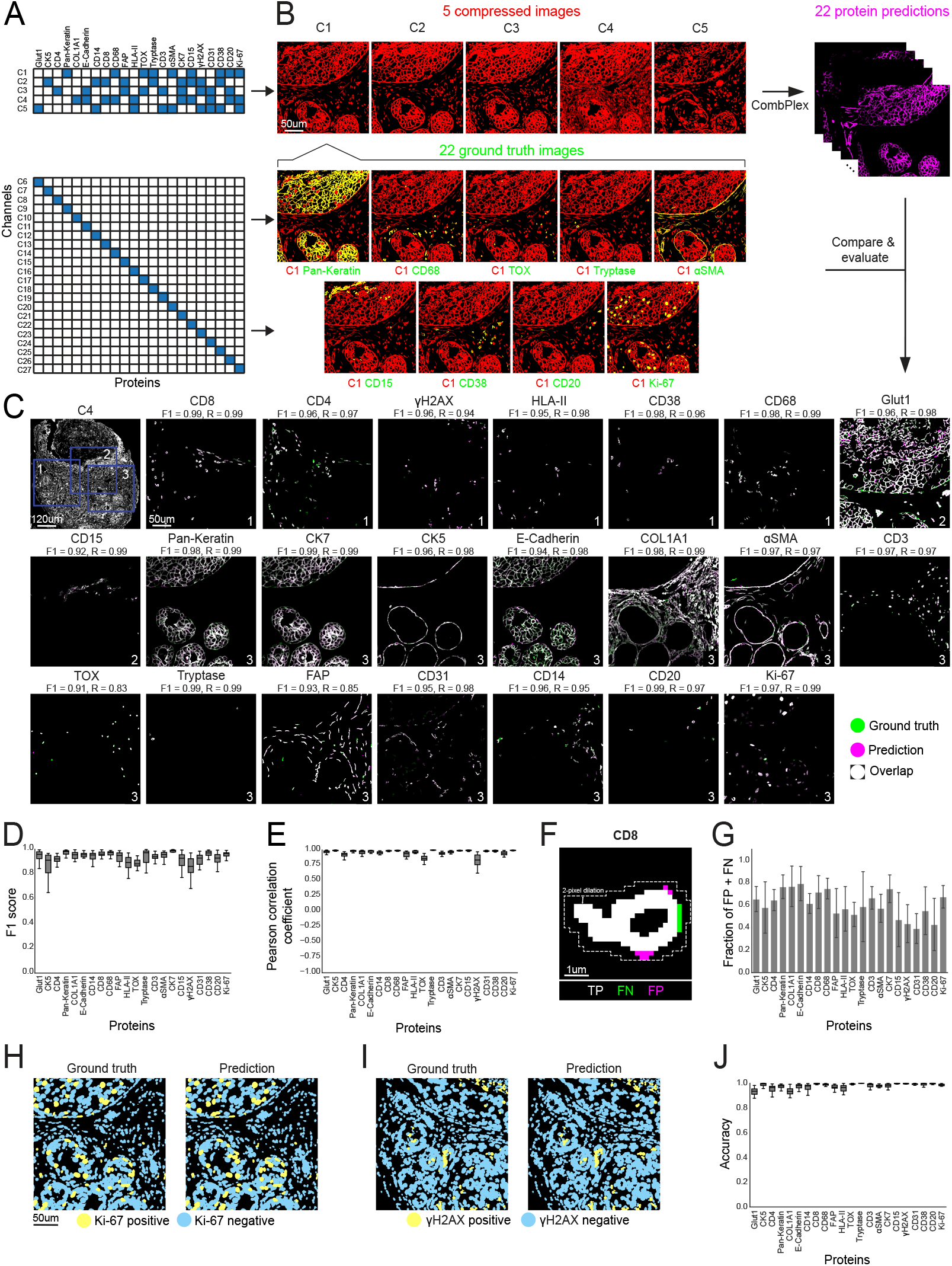
CombPlex accurately reconstructs large-scale compressions of 10 antibodies per channel. **(A)** Experimental layout. CODEX cyclic imaging was used to image 22 proteins on a single tissue section, both compressed to five measurement channels using multiple reporters per antibody (top) and individually, using a single reporter per antibody (bottom). **(B)** Top: For a single FOV, shown are the five combinatorically-compressed images (red), decompressed by CombPlex to 22 reconstructed single-protein images (magenta). Bottom: overlay of nine experimentally-measured single-protein images (ground truth, green) with a compressed image that contains all of their signals (red). Yellow pixels indicate overlap. **(C)** CombPlex reconstruction (magenta) is overlaid with the ground truth single-protein images (green). White pixels indicate overlap. For each protein image, the number in the bottom right corner indicates the location of the crop in the compressed image. **(D-E)** CombPlex was trained on experimentally-compressed images and evaluated on 30 test FOVs. For each protein (x-axis), shown are the F1 scores (D) or Pearson correlation coefficients (E) between the reconstructed and ground truth images. **(F-G)** For each protein in the 30 test FOVs, each pixel was compared between the ground truth images and the images reconstructed by CombPlex. Pixels were classified as TP (True Positives), FN (False Negatives), FP (False Positives) and TN (True Negatives). To evaluate the fraction of FP and FN pixels in the vicinity of TP signal, areas of TP signal were dilated by two pixels. The number of FP and FN pixels inside these 2-pixel dilations was divided by the total number of FP and FN pixels in the image to calculate their fraction (y-axis) for each protein (x-axis). **(H)** Cells classified as positive (yellow) and negative (cyan) for Ki-67 using ground truth (left) or reconstructed (right) images. **(I)** Same as (H) for *γ*H2AX. **(J)** Segmented cells were classified as positive or negative for each protein based on either the ground truth or the images reconstructed by CombPlex. Shown are the accuracy scores (y-axis) for each protein (x-axis).

We evaluated CombPlex’ performance in recovering the signal of the 22 individual proteins from the compressed images. We used 113 paired images for training and evaluated performance on 30 held-out images. We found that CombPlex achieved highly accurate reconstructions, both when training on real compressed images and on simulations. For the model trained on real images, median F1 scores on the test data were 0.95±0.13, Pearson correlation coefficients were 0.98±0.08, PSNR was 60.58±8.44 and SSIM was 0.998±0.018 across channels (**Fig. 4D-E, Supp. Fig. 4B-H**). Performing Pearson’s correlation only on the intersecting pixels of both images also yielded high results (R = 0.96±0.1, **Supp. Fig. 4J)**. We found that the performance of CombPlex was better on proteins with higher abundance and more robust staining (**Supp. Fig. 4I**).

Although CombPlex was overall highly accurate, we focused on the errors in prediction to better understand their properties and deduce the strengths and limitations of the approach. We separated the errors in prediction according to their type: False Negatives (FN) are pixels that contain staining for the protein in the ground truth images, but CombPlex fails to capture it, whereas False Positives (FP) are pixels in which CombPlex predicts signal that is absent in the ground truth images **(Supp. Fig. 4K)**. Visual inspection of the images suggested that FNs and FPs tended to reside on the outskirts of real signal, correctly predicted by CombPlex (True Positives, TP) **(Fig. 4F)**. To test this more rigorously, for all channels across all test images we performed a 2-pixel dilation around TP pixels **(Fig. 4F)**. We found that the majority of errors, both FN and FP, were located within a 2-pixel radius from accurately-detected signal (Fraction = 0.64±0.2, **Fig. 4G)**. Moreover, FP and FN errors were more prevalent in pixels with lower intensities and higher multiplexing level (**Supp. Fig. 5**, Methods). Taken together, we deduce that CombPlex’ errors mostly involve pixels with low intensity in the vicinity of a true signal.

Following our analysis of the errors made by CombPlex, we hypothesized that they would not materially change the expression patterns of the proteins in the cells and would have minimal downstream effects, if any, on commonly-performed tasks such as segmentation and cell classification ^29,30^. To test this explicitly, we performed whole cell segmentation with Mesmer ^31^ using nuclear and membranal proteins and found highly similar results, whether segmenting the ground truth images or the images reconstructed by CombPlex (F1 = 0.99, **Supp. Fig. 6A**,**B**). Next, we classified each protein in each cell as positive or negative and compared the cell classifications between the ground truth images and the images reconstructed by CombPlex (Methods). Overall, we found excellent agreement across all proteins (Balanced Accuracy = 0.99±0.03, **Fig. 4H-J, Supp. Fig. 6C-D**). This suggests that even in cases in which the reconstruction is not perfect, it is robust enough to enable routine downstream analyses. We conclude that CombPlex achieved accurate recovery of individual protein signals of twenty-two proteins measured using five fluorescent measurement channels.

### CombPlex accurately reconstructs compressed images measured by mass-spec imaging

Widely used multiplexed imaging approaches can be broadly grouped into two classes: fluorescence microscopy and mass-based imaging. We therefore tested CombPlex’ compatibility with Multiplexed Ion Beam Imaging by Time of Flight (MIBI-TOF), a mass-based imaging modality. Encouraged by the results of using simulations to train the network in the fluorescence data, we further evaluated whether pre-existing data, not measured in the same experiment or in the same tissue of origin, could be used to train the network. We compiled a dataset from pre-existing images in which these proteins were measured individually. We included 181 images from breast cancer, lung cancer and melanoma (**Fig. 5A**). Since the melanoma data did not contain staining for keratin, we interchanged the keratin images in this data for melan A, reasoning that both stain tumor cells and the stains are visually similar (Methods, **Supp. Fig. 7A**). The images for training were acquired in different experiments, conducted at different times, by different students, using different antibody batches and different MIBI-TOF instruments.

**Figure 5:**
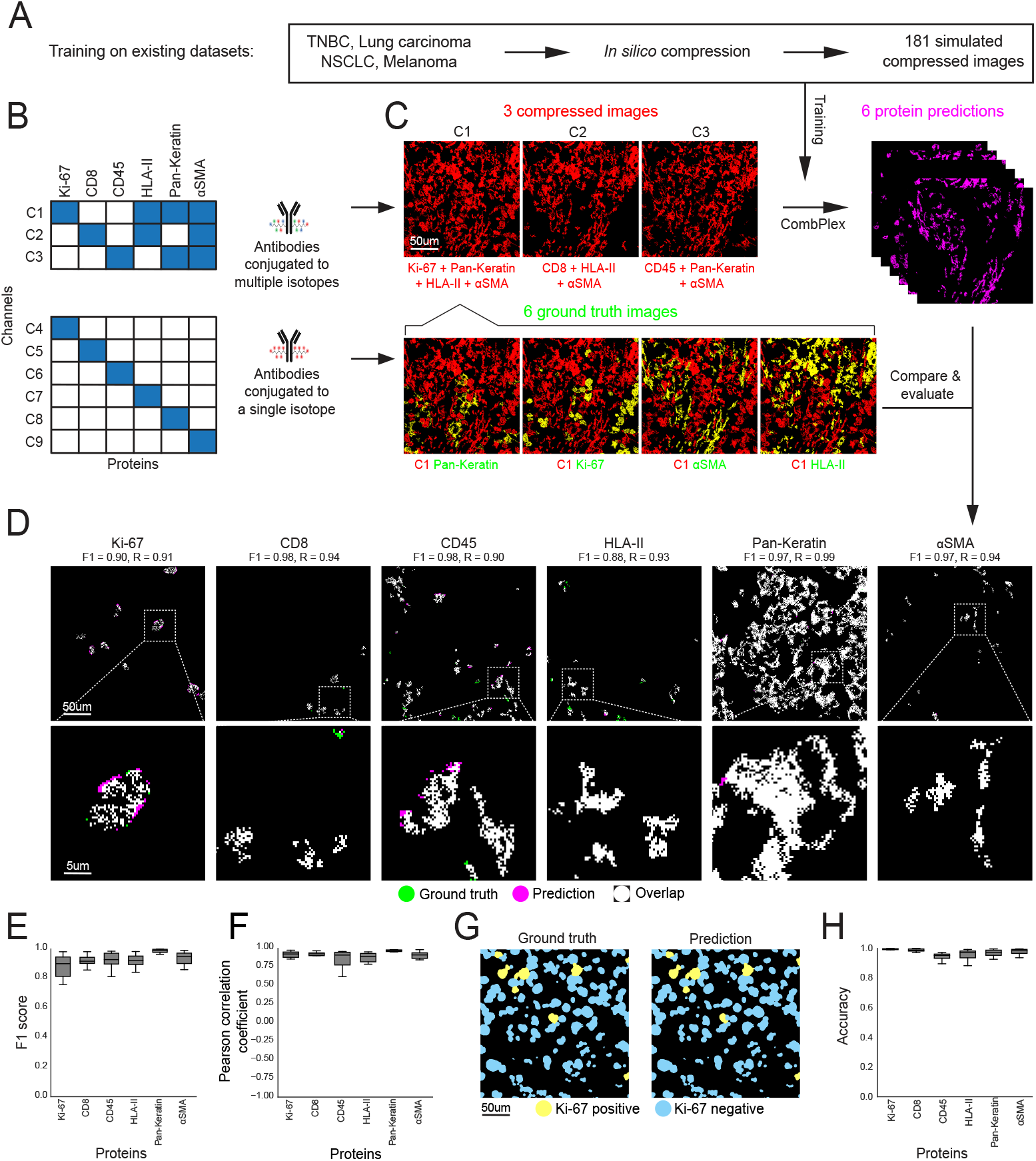
CombPlex accurately reconstructs compressed images measured by mass-spec imaging. **(A)** CombPlex was trained on compressed images simulated *in silico* from published MIBI-TOF datasets in which the proteins were imaged individually. **(B)** Experimental layout. MIBI-TOF was used to image six proteins on a single tissue section, both compressed to three measurement channels using multiple metal isotopes per antibody (top) and individually, using a single metal isotope per antibody (bottom). **(C)** Top: For a single FOV, shown are the three combinatorically-compressed images (red), decompressed by CombPlex to six reconstructed single-protein images (magenta). Bottom: overlay of four experimentally-measured single-protein images (ground truth, green) with the compressed image that contains all of their signals (red). Yellow pixels indicate overlap. **(D)** CombPlex reconstruction (magenta) is overlaid on the ground truth single-protein image (green). White pixels indicate overlap. **(E-F)** CombPlex was trained on compressed images generated by simulations from previously-available data. For each protein (x-axis), shown are the F1 scores (E) or Pearson correlation coefficients (F) between the reconstructed and ground truth images. **(G)** Cells classified as positive (yellow) and negative (cyan) for Ki-67 using ground truth (left) or reconstructed (right) images. **(H)** Segmented cells were classified as positive or negative for each protein based on either the ground truth or the images reconstructed by CombPlex. Shown are the accuracy scores comparing the results of the classification (y-axis) for each protein (x-axis).

To generate combinatorically-compressed images, we devised a method for attaching multiple metals to one antibody by loading polymers with pre-defined metal ratios and then conjugating them to the desired antibodies (methods, **Supp. Fig. 7B**). We measured on a single tissue section both the ground truth images of individual proteins and the compressed images, where the same targets were measured in a combinatorial manner (**Fig. 5B**). We stained and imaged 19 samples from both breast cancer and lung cancer. We evaluated that the compressed images indeed contained signals overlapping with the expected proteins and did not contain signals of other proteins in the panel. For example, m/z 144 contained signal matching the single-channel images of αSMA, Keratin, Ki-67 and HLA-II, but did not contain signal matching CD8 or CD45 (**Fig. 5C, Supp. Fig. 7C**). Altogether, we concluded that it is experimentally possible to multiplex several antibodies on a single measurement channel also in mass-based imaging.

We evaluated CombPlex’ performance in recovering the signal of the individual proteins from the compressed images. We used the 181 simulated-compressed images for training and evaluated performance on the experimentally-compressed images. We found that CombPlex achieved highly accurate reconstruction, with median F1 scores on the test data of 0.94±0.11 across channels, median Pearson correlation coefficients of 0.94±0.17 PSNR of 57±8 and SSIM of 0.997±0.01 (**Fig. 5D-F, Supp. Fig. 7D**,**E**). Next, we evaluated the data generated by CombPlex in cell segmentation and classification. We segmented our images using Mesmer ^31^, and classified each protein in each cell as positive or negative. We compared the cell classifications between the ground truth images and the images reconstructed by CombPlex (Methods), and found high agreement across all proteins (Balanced Accuracy = 0.97±0.01, **Fig. 5G,H, Supp. Fig. 8**). Similar to the results of the fluorescent data, this suggests that data generated by CombPlex yields cell-level classifications that are highly comparable to those obtained when proteins are measured individually.

Finally, we tested the approach in simulations on data which applied MIBI-TOF on expanded samples ^32^. The authors evaluated in their paper that the resulting images had a 3.7-fold increase in resolution, suggesting a resolution of ±140nm/pixel, which is similar to 40X. We downloaded their data and simulated combinatorically-compressed images *in silico* according to the compression matrix in supplementary figure 7F. We separated the biological specimens between the train and the test sets, using 59 images for training and 18 for testing. We found that CombPlex performed extremely well on this data (**Supp. Fig. 7G-L**, F1=0.98±0.03, R=0.99±0.02, PSNR=46±5, SSIM=0.96±0.04), suggesting its applicability also to higher resolution images.

Overall, our results indicate that CombPlex performs well also in mass-based imaging and that diverse pre-existing data can be used to train the algorithm. This greatly enhances the useability of the approach, negating the need to acquire data specifically for the purpose of training. Moreover, these results suggest that CombPlex can generalize between different tissues of origin and one model can be trained for a variety of pathologies.

### Features contributing to CombPlex performance

In CombPlex, we use a neural network to constrain the solution space of an underdetermined equation system. We explored what the network learns to aid it in its classification by performing various perturbations to the input images and training scheme. First, we evaluated whether the network learns the compression matrix *A*, since it was not directly supplied to the network during the training process. We used the published CRC CODEX dataset and the network trained on matrix *A* in figure 2A. We then tested the network’s performance on simulated compressed images that were generated by a different compression matrix *A*\* (**Fig. 6A**, Methods). We found that the network’s performance dropped by 98% compared to the original images (0.015±0.09 compared with 0.97±0.11, **Fig. 6B**). This indicates that the network indeed learns the compression scheme encoded by the matrix used for training.

**Figure 6:**
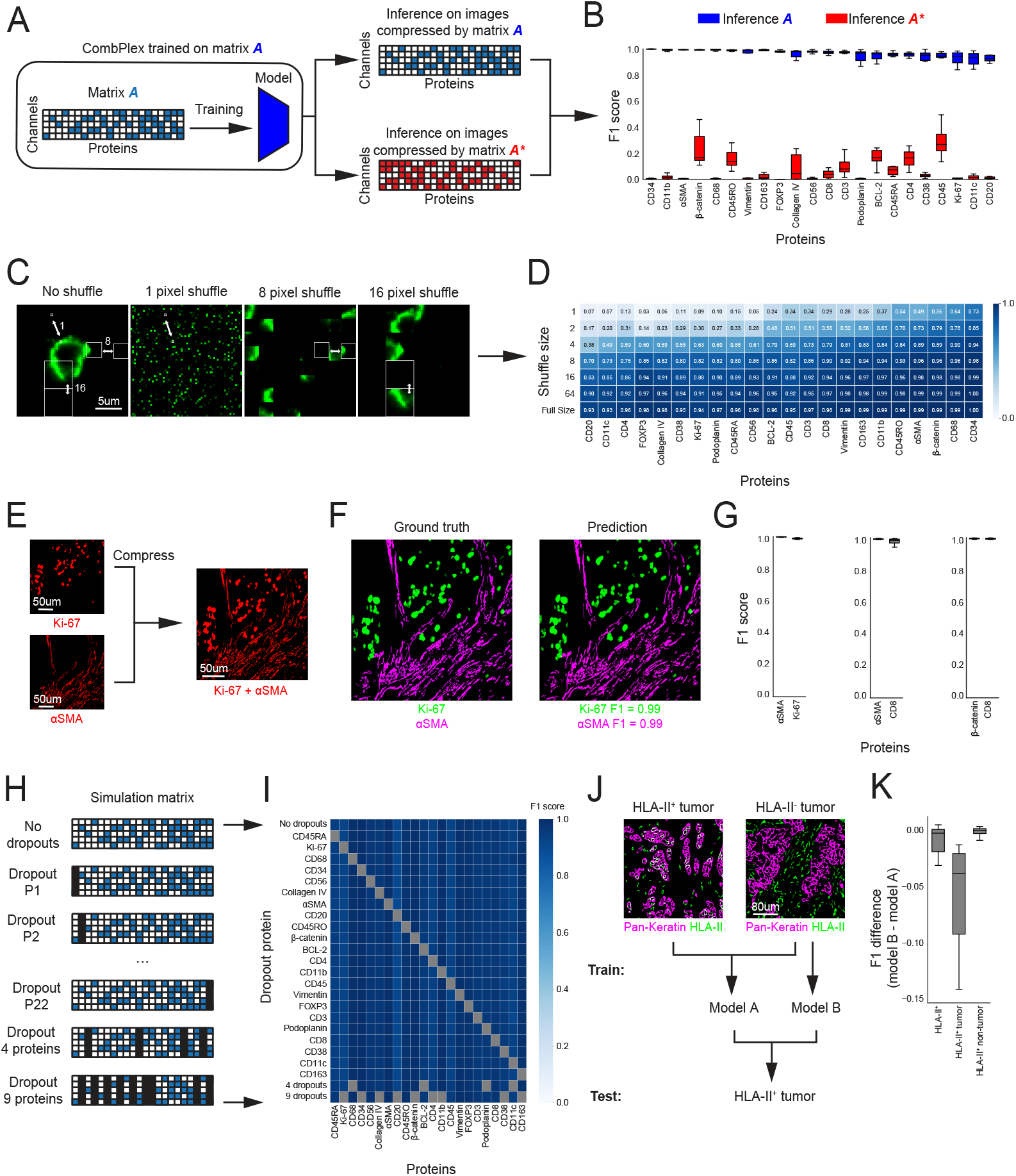
Features contributing to CombPlex performance, strengths and limitations. **(A)** For the data from Schürch et al. ^29^, a model was trained on simulated compressed images using compression matrix A. Inference was performed on simulated compressed images using compression matrices A and A*. **(B**) A comparison of F1 scores (y-axis) for each protein (x-axis) in the experiment illustrated in (A) indicates that CombPlex learns the compression matrix. **(C-D)** CombPlex was trained on the data from Schürch et al ^29^. Images of the test set were then shuffled with patches of varying sizes. (D) Shown are the median F1 scores for each protein (x-axis) on data shuffled with different patch sizes (y-axis). **(E)** Images of Ki-67 and SMA were used to simulate a naïve compression of two proteins in one image. **(F)** CombPlex was used to recover single-protein images from pairs of proteins compressed to one channel as shown in (E). Shown are the ground truth images (left) and CombPlex’ reconstruction (right). **(G)** Shown are the F1 scores (y-axis) for three 2-to-1 decompressions of pairs of proteins (x-axis) on a test set of 8 FOVs. **(H)** CombPlex was trained on the data from Schürch et al. ^29^ and then evaluated on a test set of 8 FOVs. In each run, one or more proteins were set as empty to simulate a scenario of a malfunctioning antibody or utilization of a subpanel. **(I)** The median F1 scores of different proteins (x-axis) are presented across experiments in which different proteins have been dropped out (y-axis). **(J)** Top: Shown are example images of tumor and non-tumor appearances of HLA-II (green) intersected with Pan-Keratin (marking tumor, magenta). Two models were trained on the CODEX data presented in figure (3). Model A was trained on images of HLA-II^+^ and HLA-II^-^ tumors. Model B was trained only on images of HLA-II^-^ tumors. Both models were evaluated on images of HLA-II^+^ tumors **(K)** Shown are the differences in F1 scores (y-axis) between the models in (J). HLA-II reconstruction was evaluated in the entire image (left), tumor regions (middle) and non-tumor regions (right).

Next, we explored which features of the data aided the network in converging to the correct solution. First, we evaluated the effect of spatial continuity. We took our test images from the CRC dataset and shuffled the pixels along the *x* and *y* dimensions, keeping the *z* dimension fixed. This generates images that overall have the same pixel values in each channel, but the spatial continuity of stains is disrupted (**Fig. 6C**). We found that when the size of the shuffled block is large enough to maintain single-cell morphology (>=16^2^ pixels) the network manages to produce accurate predictions (Median F1=0.96±0.12). These drop dramatically when the size of the shuffled block decreases to one pixel (Median F1=0.24±0.22), indicating that the network learns spatial continuity (**Fig. 6D**). We performed the same experiment for the MIBI-TOF data, arriving at the same conclusion also for this modality (**Supp. Fig. 9A**,**B**). Next, we tested whether CombPlex learns priors on the staining properties of different proteins. To this end, we generated a toy example in which we generated a set of compressed images of two proteins, e.g., Ki-67 and αSMA (**Fig. 6E**). These images did not contain any additional information regarding co-expression of proteins, so to decompress them back to their single-protein images, the network must learn their corresponding staining patterns. We trained CombPlex on 33 images and tested performance on 8 images. CombPlex demonstrated high recovery for both proteins (Median F1 = 0.99±0.005, **Fig. 6F,G**), indicating that it is able to learn to differentiate between the staining patterns of the different proteins.

Since CombPlex is based on supervised learning, we evaluated how robust the approach is to unexpected changes in the test data. First, we simulated an experimentally-plausible situation in which one antibody doesn’t work. In current staining schemes, if one antibody doesn’t work, only its cognate stain is affected. We were concerned that in combinatorial staining, if one antibody doesn’t work then it may ruin the reconstruction of other antibodies. To this end, we used our CRC model, trained on a compression of 22 proteins to 5 channels. We then evaluated its performance on the simulated test data, each time setting the images of one protein to zero (**Fig. 6H**). First, we examined the predictions of CombPlex for the malfunctioning protein. We found that in nearly all cases it was visually clear to a biologist that the antibody malfunctioned. Approximately half of the images were either completely empty or contained only a few isolated pixels. Most of the remaining images exhibit a fragmented appearance (**Supp. Fig. 9C**) and lower intensities than expected (**Supp. Fig. 9D**). Reassuringly, the F1 scores for the other proteins were not affected by this perturbation (Differences in F1 were 0.001±0.003 **Fig. 6I**).

Encouraged by these results, we similarly simulated scenarios in which four or nine antibodies out of the 22 were missing, and found no reduction in the F1 scores for the remaining proteins (**Fig. 6H,I**). These results highlight the robustness of the approach. Moreover, they indicate that a large model, trained on many proteins, could be used to decompress images that were combinatorically stained with only a subset of these proteins. This adds to the modularity of the approach, suggesting that a large pre-trained foundation model could accommodate the decompression of many different staining schemes visualizing different subsets of proteins.

We further tested how the approach deals with problematic antibodies, with weak or non-specific staining. We simulated weak signal of either CD8 or CD68 *in silico* by multiplying the image by a factor α, which reduced the signal to either 80%, 50% or 20% of its original level (**Supp. Fig. 9E**). We then trained CombPlex in two regimes: (A) Using the low-abundance images as training. This simulates a situation in which the target is globally weak. (B) Using the regular-intensity images as training. This simulates a situation in which the target is specifically weak in a new dataset, and this situation was not previously encountered in training. We found that in the first regime, CombPlex performed very well for both regular or reduced signal for both CD8 (median F1> 0.94, **Supp. Fig. 9F**) and CD68 (median F1> 0.96, **Supp. Fig. 9G)** even if the signal was reduced to 20% of its original intensity. We also found that for both proteins there was a slight trend to perform better with stronger signal (**Supp. Fig. 9F**,**G**). In the experiment in which regular-intensity images were used for training (regime B), we found that CombPlex tolerated well a reduction in signal intensity to either 80% or even 50%. If the signal dropped to 20%, so did the performance of the network (**Supp. Fig. 9H**,**I**). We obtained similar results in an experiment in which we simulated non-specific antibody staining (**Supp. Fig. 9J-L**). This highlights the reliance of the approach on supervised training. Unexpected behaviors, either resulting from biology or technical issues, are tolerated well if they are part of the training data. Otherwise, they could lead to a lower performance. We conclude that CombPlex needs to train on enough training data to provide good coverage of the distribution of real-world data. Our results in the MIBI-TOF experiment, trained on publicly-available data, suggest that this is feasible.

Finally, we evaluated whether the network learns rules of co-expression between proteins, and whether it can accommodate biological deviations from these expected patterns. As a test case, we used the expression of HLA-II, which is expressed on tumor cells in only a subset of patients (25% in our cohort). We took the data from Figure 3 in which we imaged seven proteins in three channels, alongside the individual stains of the seven proteins using CODEX. We then partitioned this dataset into two groups: (1) images with HLA-II+ tumor cells (N=33) and (2) images without HLA-II+ tumor cells (N=99) (**Fig. 6J**). From the first group, we took 11 images to serve as the test set. We then trained two models, designed to decompress the stains of the seven proteins. Model A was trained on 88 images from groups 1 and 2. Model B was trained on 88 images only from group 2 (**Fig. 6J**). Consequently, during training model A had the chance to learn that HLA-II and Pan-Keratin could be coexpressed, whereas model B had the chance to learn that they are not coexpressed. We then used both models to decompress the test set. We found that both models were equally successful at reconstructing HLA-II on antigen-presenting cells, but there was a 2-9% reduction in performance in reconstructing HLA-II expression on tumor cells when such cells were not presented at all in the training data (**Fig. 6K**). We conclude that CombPlex can generalize to novel biology beyond the training set, but these capabilities are limited and, like in any supervised learning approach, ideally the training set should cover the full diversity of the domain.

Overall, we demonstrated that the network learns the compression matrix, spatial continuity of signals, the staining patterns of proteins and their co-expression, and leverages these properties in order to decompress combinatorically-multiplexed images. It can easily generalize to accommodate missing channels, but care should be taken with biological deviations that are not present in the training set.

## Discussion

Current approaches to image multiple proteins *in situ* are inherently limited in throughput by their design: using a separate measurement channel for each individual protein. Combinatorial staining holds great potential, because it exponentially increases the number of proteins that can be measured using any imaging platform from *C* to a theoretical limit of 2^*c*^ − 1. For example, using three-color fluorescent microscopy one could measure seven proteins (2^3^ − 1). Seven-color imagers, well-established in research and in the clinic, could serve to image dozens of proteins, and a multiplexed imaging instrument could accommodate hundreds of targets. However, to date, combinatorial staining was not applied to proteins, presumably due to the high abundance of proteins in cells. High protein density makes it impossible to resolve the signals of individual proteins under standard magnification conditions and thus mathematically infeasible to reconstruct the individual protein expression images.

Here, we presented CombPlex, an experimental approach to perform combinatorial staining of proteins, coupled with an algorithmic framework to accurately decompress the combined signal back to individual protein images. We showed that CombPlex allows us to decouple the staining of individual proteins stained together with over 90% accuracy, even in low, 20X magnification, both in fluorescent and in mass-based imaging. We showed that although the problem of reconstructing individual protein images from compressed measurements is mathematically underdetermined, CombPlex manages to achieve highly accurate reconstructions. Our analyses suggest that it does so by leveraging various features of the data, including spatial continuity, individual protein expression patterns and protein co-expression. While it is infeasible to encode such features explicitly, a neural network trained on image examples can learn them and exploit them as strong priors for probable solutions for solving the reconstruction task.

Combinatorial staining poses several challenges, both experimental and algorithmic. It remains to be determined how many proteins can be multiplexed on a single channel without signal deterioration, devising methods to quickly titrate antibodies such that they could be imaged with similar exposure times, determining which antibodies could be multiplexed, reducing the cost of antibodies etc. Our experiments indicated that at least 10 antibodies could be combined on a single channel. CombPlex performs well for many antibodies, spanning a range of signal strengths, and that the performance of CombPlex improves with more robust staining. However, if one is interested in a target which has very weak or has strong non-specific staining, one should take care in multiplexing it.

Solving the decompression problem is a non-trivial algorithmic challenge. CombPlex solves it utilizing supervised deep learning to predict both the binary mask and the values for each protein. This design choice has led, in our hands, to the best performance. However, it also has some drawbacks. Predicting the binary mask requires clean images, which are time-consuming to generate. Utilizing supervised deep learning means that the network is trained on a set of tissues, on a pre-defined set of proteins. Deviations from the training set, either due to technical or biological reasons, could lead to reduced performance. Here we showed that CombPlex was able to generalize across several cancer types and accommodate reasonable technical malfunctions, but further exploration of failure cases is warranted. Such issues could be overcome by training on large datasets that encompass the full technical and biological variability of real-life data. Future algorithmic developments could explore strategies for intelligent design of the compression matrix, circumventing the need for the binary network or developing non-supervised or self-supervised approaches.

CombPlex requires training data in the form of individual protein images, coupled with their combinatorically-compressed counterparts. In the manuscript, we presented several results that could reduce the entry barrier posed by the need to acquire such training data. We devised a method to simulate combinatorically-compressed images *in silico* from individual measurements of the corresponding proteins. We demonstrated that a network trained on simulated images achieves high performance, almost the same as a network that was trained on experimental combinatorically-compressed images. Using simulations to train the network holds great advantages. It negates the need to acquire both single-protein images and combinatorically-compressed images on the same tissue. Here, we showed that we could train CombPlex on diverse, pre-existing data from various tissue sources and use this network to generate accurate reconstructions of experimentally compressed data. Moreover, we showed in simulations that CombPlex could accommodate 30% missing proteins without a reduction in performance for the remaining proteins. Putting these results together, we envision that a community resource could be generated in which hundreds of proteins would be measured individually on various tissues using currently available multiplexed imaging approaches, such as cyclic fluorescence or mass-based imaging. This resource could then serve as the basis to pretrain a large foundation model that accommodates the decompression of various staining schemes, visualizing different subsets of proteins. Subsequent studies could then use CombPlex to image multiple proteins on their respective samples, saving valuable time and money. Notably, such community efforts are well on their way, examples including the Human Protein Atlas, Single Cell Human Atlas, or the Human BioMolecular Atlas Program ^33–35^.

Protein imaging is a pillar of pathology. While recent technological developments have increased the number of proteins visualized *in situ*, these increases have been linear and thus far less than required for proteome-scale measurements. Looking forward, it is difficult to envision how proteome-wide measurements could be performed *in situ* without the use of combinatorics. As such, this study lays a foundation for a multitude of experimental, analytical and biological endeavors to undoubtedly follow. Farther-reaching projects could include applying CombPlex to measurements of abundant mRNAs that currently challenge the spot-counting requirements of barcoded *in situ* hybridization methods. We could also expect to see further increases in throughput using standardized combinatorically-multiplexed antibody panels, e.g., a standard immune profiling kit. These could be integrated with available instruments for multiplexed imaging to further increase their multiplexing capabilities. Finally, it will be enticing to observe further experimental and algorithmic developments that could reduce the amount of training data needed, simplify its acquisition or eliminate it altogether.

## Methods

### CombPlex overview

CombPlex consists of two components: 1) an experimental approach for combinatorial staining of proteins, and 2) an algorithmic framework for decompression of combinatorically-compressed images.

The experimental approach aims to generate compressed multiplexed images, where each channel contains the signal of multiple proteins. To generate these experimentally, multiple proteins are measured simultaneously on the same channel, according to a given compression matrix. Calibration of this staining regime should be optimized according to these principles:

- Protein signals in each channel must be visible under the same imaging conditions.
- A protein that is measured in more than one channel should display similar staining patterns across channels.

The algorithmic approach is divided into two steps. The first aims to estimate the regions where individual protein signal exists, while the latter completes the recovery by estimating pixel intensities in those regions. The input for CombPlex is a multiplexed image denoted as *y* ∈ *R*^*H*×*W*×*C*^, which is a compressed representation of the underlying signal *x* ∈ *R*^*H*×*W*×*P*^ (where *P* > *C*). The output is a reconstructed signal *x*′ of the underlying signal *x*.

- *H* and *W* are the height and width dimensions, respectively.
- *C* signifies the number of channels in the measurement (e.g., colors captured in a microscope).
- *P* denotes the total number of distinct underlying proteins that were subjected to imaging during the experimental procedure.

An illustration of the approach is shown in Figure 2G and Supplementary figure 1.

### Algorithm

#### Generating the compression matrix

The compression scheme is determined by a binary matrix *A* with *C* rows and *P* columns, representing the measurement channels and individual proteins, respectively. In this matrix, a value of 1 at position (*i, j*) indicates that protein *j* is stained in measurement channel *i*. To generate matrix *A* we employ three principles, ordered by their priority:

1. Each column vector must be unique such that different proteins are encoded differently.
2. The sum of each column is minimized to facilitate the process of partitioning the antibodies.
3. The sums of the rows are balanced, which aids in the computational task of distinguishing a higher number of protein signals within the same compressed image.

In the GitHub repository associated with the project, we supply a script to assist in constructing this matrix: https://github.com/KerenLab/CombPlex. We note that if one uses the full degrees of freedom of compression, as we did for example in our experiments compressing seven proteins to three channels, then steps 2 and 3 are unnecessary. In these cases, all possible vector combinations containing 0 and 1 will be used.

After constructing the matrix, the next task is to assign the different proteins to the *P* columns. For our CombPlex experiments presented in Figures 2-6, we chose the assignment of proteins to the columns empirically following simulations. We first collected the individual images (e.g. of the 22 proteins) in isolation. We then simulated compressed data using 8 random compression matrices and picked the best-performing matrix to generate experimentally-compressed data in the following experiments. We could not exhaustively search the entire space of optional compressions (22! options). It is possible that there is a compression matrix that will outperform the one that we ultimately chose. We foresee that future research will add valuable insight into better approaches for selecting the compression scheme to maximize performance.

#### Simulations of combinatorically-compressed images

Given single-protein images *x*, we simulated compressed images *y* (**Supp. Fig. 1A**). We conducted a comparison between experimentally compressed images and those generated by summing intensities over corresponding single-protein images. While the comparisons revealed high correlations, there were still noticeable levels of noise and skewness, as illustrated in supplementary figure 3F. Therefore, our simulation procedure adheres to the fundamental concept of linearly adding the single-protein images, along with different noise sources and augmentations. Simulations for MIBI and CODEX were performed differently due to the distinct characteristics of the signal in each technology.

CODEX simulation process:

1. Noise was applied on Matrix *A*, followed by convolution with the single-protein images *x*:

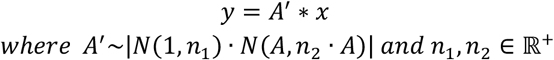 This noise accounts for experimental differences between channels, resulting from either systematic differences in channel intensities or stochastic differences in antibody titers.
2. Pixel-wise noise was applied on *y* to account for pixel-wise errors during imaging and accumulated background during simulation:

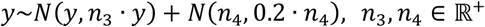

MIBI-TOF simulation process:

1. Noise applied on Matrix A as described in the CODEX simulation process, creating *A*^′^.
2. Convolving the single-protein images with Matrix A’:

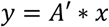
3. The compressed images were split to 2×2 patches, and each patch was randomly rotated.

#### Reconstructing individual protein images

During our explorations of the data, we found that from a biologist’s point of view, false predictions of signals are often more crucial than the precise reconstruction of abundant signals. As such, there is a need to reconcile between two incentives: accurately reconstructing the location of the signals and their magnitude. Therefore, CombPlex is designed as a combination of two CNNs: a masking network, and a value reconstruction network. The goal of the former is to determine *where* different proteins are expressed in the image, while the goal of the latter is to determine *how much* each protein is expressed in each pixel of the image.

The input to the masking network is a compressed image with *C* channels (*y*), and the output is *P* images with probability values 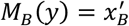. The probability value in a pixel in image *p* indicates the likelihood of the pixel to contain a signal of protein *p*. To train the network we perform supervised learning using pairs of compressed images (*y*) and their corresponding individual-protein images (*x*). The goal is to minimize the binary cross-entropy between the binarized GT *x*_*B*_ and *M*_*B*_(*y*). In parallel, we train the value reconstruction network. The input is again a compressed image with *C* channels (*y*), but the output is *P* images with predictions for the intensity values 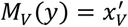. We train it similarly in a supervised manner, to minimize the MSE between the GT single-protein images *x* and *M*_V_(y).

To merge these outputs, a threshold of 0.5 is applied to the probability mask. For each pixel with a probability higher than 0.5, its predicted value is extracted from the output of the value reconstruction network using pixel-wise multiplication:

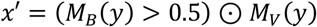

Training the decompression masking network *M*_*B*_

The goal of *M*_*B*_ is to recover the binary support of the individual protein images (**Supp. Fig. 1B**). We perform pixel-wise classification for each pixel (*i, j*) in the image with respect to the presence or absence of each of the *P* proteins. Mathematically, we recover a function denoted as *M*_*B*_ (*y*), which takes a compressed image *y* ∈ *R*^*H*×*W*×*C*^ as input and outputs a probability map 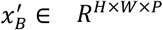 such that:

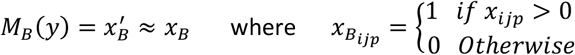

The network is trained in a supervised manner using pairs 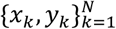 of compressed images and their corresponding individual protein images. The compressed images can be either acquired experimentally or simulated as described in the *Simulations of combinatorically-compressed images* section.

The objective is to minimize the binary cross-entropy (BCE) loss between the real *x*_*B*_ and the model output *M*_*B*_(*y*) in a set of *N* images. Formally, finding a set of parameters *θ* to minimize:

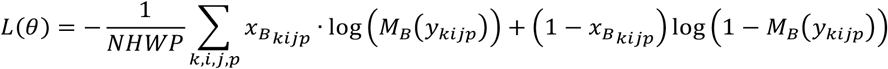

The network follows the UNet 3+ architecture, an extension of the original UNet architecture that introduces full scale skip connections and convolutional blocks ^36^. The symmetric encoder-decoder structure cosists of five scales in terms of depth, with convolutional layers having a kernel size of 3×3. Batch normalization layers and ReLU activation functions are applied after each convolutional layer. We used Adam optimizer, constant learning rate value of 0.0001 and batches of 4 FOVs per iteration. Data augmentations were used to enrich the datasets: random crop of size 512×512, random flips and rotations by 90 degrees.

##### Training the value reconstruction network *M*_*V*_

The goal of *M*_*V*_ is to recover the intensities of the individual protein images (**Supp. Fig. 1B**). We perform value reconstruction for each pixel (*i, j*) in the image for each of the *P* proteins. Mathematically, we recover a function denoted as *M*_*V*_(y), which takes a compressed image *y* ∈ *R*^*H*×*W*×*C*^ as input and outputs a set of single-protein images 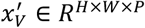.

The training methodology for this network closely follows the approach detailed for the decompression masking network, except for two key modifications. Firstly, the sigmoid activation layer after the model architecture is omitted, enabling the prediction of values beyond one. Second, the training process adopts different loss functions to for reconstructing the values according to the nature of the data. For CODEX data, we use MSE loss:

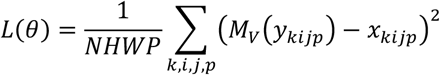

For MIBI-TOF data, which is a stochastic counting process, we employ the Poisson negative log likelihood loss:

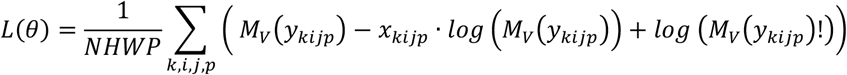

While *log*(*M*_*V*_(*y*_kijp_)!) is approximated using the Stirling approximation term:

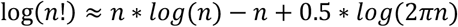

##### Model ensembling

To enhance stability and performance, we utilize an ensemble of models for some of the experiments. Suppose we have trained decompression masking networks, denoted as 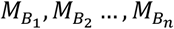, and trained value reconstruction networks, denoted as 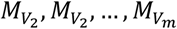. To create an ensemble, we establish a threshold *T* and combine the models by averaging their predictions.

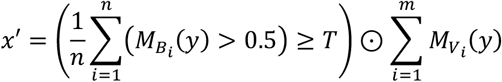

### CODEX *in silico* experiment

#### Data

Analysis was performed on data from Schürch et al. ^29^, in which 57 proteins were measured in colorectal cancer (CRC) specimens using co-detection by indexing (CODEX) cyclic fluorescence ^29^. From the publicly available dataset, 22 proteins which had specific staining and a high signal-to-noise ratio, were chosen for compression (**Supp. Table 1A**).

#### Image pre-processing

Multiplexed image sets from 41 FOVs were initially preprocessed as described in Schurch et al ^29^ to perform cycle registration and autofluorescence removal (**Supp. Fig. 10A**). Subsequently, images were denoised using the Ilastik software, by training a pixel classifier for each protein ^37^. Briefly, all FOVs for a given protein were loaded into Ilastik and used for training the pixel classifier by manually marking real signal and noise. The trained model was used to produce binary segmentation masks for each FOV. Raw images were intersected with the segmentation masks to produce clean images, containing the original pixel values where the segmentation mask pixels equaled 1. Images were filtered for antibody aggregates by applying average smoothing with a sliding window and removing signal clusters with a radius smaller than 100 pixels (**Supp. Fig. 10B, Supp. Table 1A**).

#### Simulation and decompression

The compressed images *y* were simulated using the single-protein images *x* according to: *y* = *A* ∗ *x*. Matrix *A* was generated as described in the *“Generating the compression matrix”* methods section. To train both masking and value reconstruction models, the dataset was split into a training set of 33 FOVs and a test set of 8 FOVs. The models were trained with noise parameters *n*_1_ = *n*_2_ = *n*_4_ = 0 and *n*_3_ = 0.01 for 316K and 586K batches accordingly, using a GPU of type Nvidia Quadro RTX8000.

### CODEX compression experiment

#### Antibody conjugation

A summary of antibodies, barcode oligos, reporter fluorophores and concentrations can be found in Supp. Table 1B-C. For the small-scale experiment, oligo-conjugated primary antibodies were prepared 50µg at a time using the Akoya Antibody Conjugation Kit (Akoya Biosciences, Delaware, USA) according to the manufacturer’s recommended protocol. Biotin-conjugated antibodies were prepared 100µg at a time using the ChromaLINK One-Shot Antibody Biotinylation Kit (Vector labs, Newark, USA). For the large-scale experiment, oligo-conjugated primary antibodies were prepared 50ug at a time precisely as detailed in Black et al. ^38^.

#### CODEX staining

Breast cancer TMA glass slides were acquired from TissueArray (Maryland, USA). Slide tissue sections were baked at 70°C for 20 min. Tissue sections were deparaffinized with 3 washes of fresh xylene. Tissue sections were then rehydrated with successive washes of ethanol 100% (2x), 95% (2x), 80% (1x), 70% (1x), and ultra-pure DNase/RNase-free water (Bio-Lab, Jerusalem, Israel), (3x). Washes were performed using a Leica ST4020 Linear Stainer (Leica Bio-systems, Wetzlar, Germany) programmed to 3 dips per wash for 30 seconds each. The sections were then immersed in epitope retrieval buffer (Target Retrieval Solution, pH 9, DAKO Agilent, Santa Clara, CA) and incubated at 97°C for 40 min and cooled down to 65°C using Lab vision PT module (Thermofisher Scientific, Waltham, MA). For the small-scale experiment, slides were stained using Akoya Staining Kit for Phenocycler (Akoya Biosciences, Delaware, USA) according to the manufacturer’s recommended protocol. For the large-scale experiment, slides were stained as detailed in Black et al. ^38^. Details on the staining conditions for the primary antibodies are provided in Supp. Table 1C.

#### CODEX imaging

CODEX imaging was performed using a Phenocycler Fusion system (Akoya Biosciences, Delaware, USA) with an Olympus 20X/0.7 imaging objective. Phenocycler reporter plate and Flow Cell assembly were performed according to the manufacturer’s protocol. Compressed images were created by hybridizing multiple reporters loaded with the same fluorophore in the same imaging cycle. Details on the layout of the reporter plate and imaging conditions are provided in Supp. Table 1C. For each antibody, the working titer was determined as the titer resulting in the detection of all antibodies in each compressed channel. Titers were assessed by overlaying compressed and ground truth images and evaluating successful compression both qualitatively and quantitatively by measuring signal correlation (**Supp. Fig. 3D-F**).

#### Image pre-processing

Multiplexed images underwent pre-processing within the Phenocycler, as implemented by the manufacturer. Images were registered over imaging cycles and background autofluorescence was removed by subtracting an image of a blank channel (**Supp. Fig. 10A**). Multiplexed image sets were extracted by manually marking individual cores using QuPaths’ TMA De-arrayer with a diameter of 1.2mm and a custom groovy script, which exports them as tiff files ^39^. Single-protein images were denoised and aggregate-filtered as described above (**Supp. Fig. 10B, Supp. Table 1B**). To denoise the combinatorically-compressed images, compressed segmentation masks were produced by a binary summation of all single-protein images in each compressed channel, according to the compression matrix. Experimental combinatorically-compressed images were intersected with the compressed segmentation mask to produce clean compressed images.

#### Dataset and model settings

The dataset for the small-scale experiment included 132 1mm^2^ FOVs and was split into a training set of 100 FOVs and a test set of 32 FOVs. The dataset for the large-scale experiment included 143 1mm^2^ FOVs and was split into a training set of 113 FOVs and a test set of 30 FOVs.

For each of the experiments (small and large scale) we trained four types of models: a masking model and a reconstruction model trained on experimentally compressed images, and a masking model and a reconstruction model trained on simulated compressed images as described in the *Simulations of combinatorically-compressed images* section. All models were evaluated on the experimentally compressed images.

In the small-scale experiment trained on experimentally compressed images, the binary model was trained for 159K batches and the reconstruction model was trained for 602K batches. Both models were trained with noise parameters *n*_1_ = *n*_2_ = *n*_4_ = 0 and *n*_3_ = 0.01. In the small-scale experiment trained on simulated compressed images, the binary model was trained with noise parameters *n*_1_ = *n*_4_ = 0, *n*_2_ = 0.4, *n*_3_ = 0.35 for 150K batches and the reconstruction model was trained with noise parameters *n*_1_ = *n*_4_ = 0, *n*_2_ = 0.2, *n*_3_ = 0.35 for 418K batches.

In the large-scale experiment trained on experimentally compressed images we employed an ensemble of models for the masking and reconstruction. Each ensemble was comprised of 7 models trained with noise parameters *n*_1_ = *n*_2_ = *n*_4_ = 0 and *n*_3_ ∈ [0.01,0.1], using threshold *T* = 0.5. Each masking model was trained between 200K to 400K batches and each reconstruction model was trained between 400K to 600K batches. In the large-scale experiment trained on simulated compressed images, we employed an ensemble of models for the masking and reconstruction. Each ensemble was comprised of 4 models. The binary models were trained with noise parameters *n*_1_ = 0.2, *n*_2_ = 0.4, *n*_3_ = 0.35 for 430K-530K batches using threshold *T* = 0.75. The reconstruction models were trained with noise parameters *n*_1_ = 0, *n*_2_ ∈ [0.1,0.4], *n*_3_∈ [0.1,0.5], *n*_4_ ∈ {0,500} for 300K-530K batches.

All models were trained using GPUs of type NVIDIA A100-SXM4-80GB and Quadro RTX8000.

#### Cell classification

Cell segmentation masks were generated using Mesmer and the nuclear channel (DAPI) for each FOV ^31^. Intensity values were transformed using arcsinh transformation with a factor of 0.1 and normalized to 0-1 using min-max normalization between the 1^st^ and 99.5^th^ percentiles. Thresholds for gating positive/negative were chosen for each protein in the ground truth images using visual inspection. Optimal thresholds for the reconstructed images were chosen using a ROC curve analysis, which maximized accuracy between the classification in the ground truth and reconstructed images.

### MIBI-TOF experiment

#### TMA preparation for MIBI-TOF

Samples of ten breast carcinoma and seven lung carcinoma FFPE biopsies from ten patients were obtained from Hadassah Hospital Pathology department and Sheeba Hospital respectively. IHC staining for tumor cells (Pan-Keratin) and immune cells (CD45) was performed on all samples as described below. IHC results were reviewed for expression of both proteins and 2mm cores were compiled into a TMA.

#### Antibody conjugation

A summary of antibodies, reporter isotopes, and concentrations can be found in Supp. Table 1D. Metal conjugated primary antibodies were prepared 100ug at a time using the Maxpar X8 Antibody Labeling Kit (Fluidigm, California, USA) according to the manufacturer’s recommended protocol. Multi-metal conjugates were prepared using the same kit and protocol, by mixing multiple isotopes to a final volume of 5µl and conjugating them to a single polymer, as detailed in Supp. Table 1E and supplementary figure 7B For each antibody, the ratio between isotopes was calculated to compensate for inherent differences in their respective ionization efficiencies (I/E) ^15^ as listed in table S1E. Following metal labeling, antibodies were diluted in Antibody Stabilizer PBS solution (Boca Scientific, Washington, USA) to 0.5 mg/ml and stored long-term at 4°C.

#### MIBI-TOF staining

Staining was performed as detailed in Elhanani et al ^40^. Tissue sections (5µm thick) were cut from FFPE tissue blocks of the TMA using a microtome, and mounted on gold-coated slides (Ionpath, Menlo Park, CA) for MIBI-TOF analysis. Slide tissue sections were baked at 70°C for 20 min. Tissue sections were deparaffinized with 3 washes of fresh xylene. Tissue sections were then rehydrated with successive washes of ethanol 100% (2x), 95% (2x), 80% (1x), 70% (1x), and ultra-pure DNase/RNase-Free water (Bio-Lab, Jerusalem, Israel), (3x). Washes were performed using a Leica ST4020 Linear Stainer (Leica Bio-systems, Wetzlar, Germany) programmed to 3 dips per wash for 30 s each. The sections were then immersed in epitope retrieval buffer (Target Retrieval Solution, pH 9, DAKO Agilent, Santa Clara, CA) and incubated at 97°C for 40 min and cooled down to 65°C using Lab vision PT module (Thermofisher Scientific, Waltham, MA). Slides were washed for 5 min with a wash buffer made with ultra-pure DNase/RNase-Free water (Bio-Lab, Jerusalem, Israel) and 5% 20X TBS-T pH7.6 (Ionpath, Melo Park, CA). Sections were then blocked for 1h with 3% (v/v) donkey serum (Sigma-Aldrich, St Louis, MO) diluted in TBS-T wash buffer. Metal-conjugated antibody mix was prepared in 3% (v/v) donkey serum TBS-T wash buffer and filtered using centrifugal filter, 0.1 µm PVDF membrane (Ultrafree-Mc, Merck Millipore, Tullagreen Carrigtowhill, Ireland). Two panels of antibody mixes were prepared. The first panel contained most of the metal-conjugated antibodies, and was incubated overnight at 4°C in a humid chamber. The second mix contained antibodies for dsDNA and αSMA. Following overnight incubation, slides were washed twice for 5 min in wash buffer and incubated with the second antibody mix for 1h at 4°C. Slides were washed twice for 5 min in wash buffer and fixed for 5 min in diluted glutaraldehyde solution 2% (Electron Microscopy Sciences, Hatfield, PA) in PBS-low barium (Ionpath, Menlo Park, CA). Tissue sections were dehydrated with successive washes of Tris 0.1 M (pH 8.5), (3x), Ultra-pure DNase/RNase-Free water (2x), and ethanol 70% (1x), 80% (1x), 95% (2x), 100% (2x). Slides were immediately dried in a vacuum chamber for at least 1 h prior to imaging.

#### MIBI-TOF imaging

Imaging was performed using a MIBIScope mass spectrometer (Ionpath, Menlo Park, CA) with a Xenon ion source. FOVs of 400µm^2^ were acquired using a grid of 1024×1024 pixels, with 1msec dwell time per pixel. For each antibody, the working titer was determined as the titer that resulted in the detection of all antibodies in each compressed channel. This was assessed by overlaying compressed and ground truth images and evaluating successful compression (**Supp Fig. 7C**).

#### Image pre-processing

Training data for MIBI was collected from published datasets, as well as unpublished data in the lab. For published data, we used the clean images provided by the authors. For new data, single-protein images were extracted, slide background-subtracted, denoised and aggregate filtered using MAUI ^41^ (**Supp. Fig. 10B**). Briefly, MAUI uses a k-nearest-neighbor approach to generate a histogram of signal density in the image. A biologist then applies a threshold over this histogram. Regions with a signal density below the threshold are classified as background and are zeroed. Following this step, we perform “aggregate-removal”. Metal-conjugated antibodies may sometimes form dense aggregates, which are present in the images as small specs with abnormally high values. We remove these by applying a Gaussian filter to the image and removing all connected component < 100 pixels. This process was performed for both ground truth single-protein measurements as well as experimentally-compressed combinatorial images. To denoise the combinatorically-compressed images, compressed segmentation masks were produced by a binary summation of all single-protein images in each compressed channel, according to the compression matrix. Experimental combinatorically-compressed images were intersected with the compressed segmentation mask to produce clean combinatorically-compressed images.

#### Dataset and model settings

The decompression masking network was trained and tested using MIBI-TOF images of compressed or single expression measurements of Pan-Keratin, αSMA, CD45, CD8, HLA-II, and Ki-67 from several datasets:

- *Triple-negative breast cancer dataset:* The dataset contains 44 800×800 µm^2^ images ^42^.
- *Non-small cell lung carcinoma dataset:* The dataset contains 84 800×800 µm^2^ images.
- *Melanoma lymph node dataset:* The dataset contains 15 800×800 µm^2^ images.
- *Breast and Lung Carcinoma TMA dataset:* The dataset contains 57 400×400 µm^2^ images.

The dataset included 200 FOVs of varying sizes and was split into a training set of 181 FOVs and a test set of 19 FOVs. Training was performed using simulated compressed images as described in the algorithm section. Inference was performed on experimental combinatorically-compressed images. We employed an ensemble of models for the binary mask output, comprised of 3 models with noise parameters *n*_1_ = 0, *n*_2_ ∈ [0.25,0.4] and threshold *T* = 1, and trained for about 20K batches each. The value reconstruction model was trained with noise parameters *n*_1_ = 0, *n*_2_ = 0.4 for 120K batches. All training procedures were using a GPU of type Nvidia Quadro RTX8000.

#### Cell classification

Cell classification for the MIBI-TOF experiment was performed as described above.

#### MIBI-TOF expanded samples

Analysis was performed on data from Bai et al ^32^. The dataset included 77 400um^2^ FOVs and was split into a training set of 59 FOVs and a test set of 18 FOVs. Training was performed using simulated compressed images as described in the algorithm section. The compression matrix was generated as described in the *“Generating the compression matrix”* methods section. The binary model was trained for 46K batches and the reconstruction model was trained for 70K batches. Both models were trained with noise parameters *n*_1_ = *n*_2_ = *n*_4_ = 0 and *n*_3_ = 0.01, using a GPU of type Nvidia Quadro RTX8000.

### Analyses

#### Compressed sensing

Gradient descent was performed to minimize the objective function, which consists of three components:

- MSE loss between the reconstructed compressed images, created by convolution of single protein predictions with *A*, and the simulated compressed images.
- Sparsity constraint on the single-protein reconstructed images, factored by *α* = 0.1.
- Smoothing constraint on the single-protein reconstructed images, factored by *β* = 0.001.

Mathematically, we sought to find 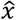 that minimizes:

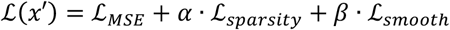

Where:

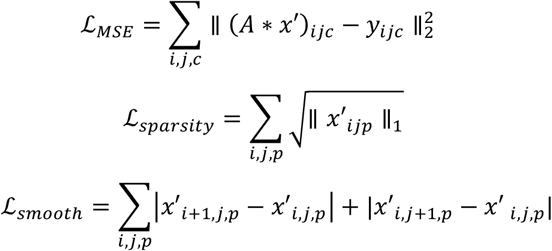

#### Decompression using a one-step algorithm

The CombPlex algorithmic framework was replaced by a single value reconstruction network, using MSE loss. The dataset, the process of simulating compressed images, the architecture of the model, and the hyperparameters employed were as outlined in the CODEX in-silico experiment. The results were noisy and to better separate between signal to noise, a threshold of 500 was applied across all proteins and FOVs.

### Considerations for experimental design

In CombPlex, multiple antibodies are imaged together, and then separated computationally using a pre-trained neural network. Below we indicate some experimental considerations in choosing reagents and multiplexing them:

1. For CombPlex, it is best to work with robust reagents – strong antibodies with low background and non-specific staining.
2. Antibodies should be tittered such that they can be imaged using a fixed exposure.
3. The training set should ideally cover the distribution of real-world data in terms of antibody intensities, staining patterns etc.

### Evaluation metrics

In the manuscript we used several metrics to evaluate the decompression. We discuss their calculation, as well as their properties, strengths and weaknesses below.

*F1*

F1 scores were computed by comparing the GT single-protein images with their recovered image using CombPlex. This was done separately for each FOV and each protein using the following formula:

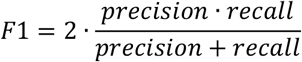

Since MIBI data is pixelated, prior to calculating the F1 score, we applied an average filter with a kernel size of 5×5 pixels (2×2μm^2^) on the GT single-protein images to smooth the signal and intersect each of them with the compressed images containing the specific protein.

We found that pathologists and biologists are often most interested in recovering the presence or absence of staining, as well as maintaining the overall interrelations of image intensities during reconstruction, rather than focusing on the exact intensity values. Precise values in such images may often vary based on technical or experimental variations, such as differences between antibody batches, the brightness of the fluorophore used to detect the antibody or slight changes in detector sensitivity. As such, the F1, which measures the degree of overlap of two binary signals, is well-suited for estimating this presence or absence of staining. It is also particularly well-suited for sparse data. Many of the protein images could contain just a few positive cells. In such cases, correlation, PSNR and SSIM will show artificially good results, since most of the image is negative for the signal. In contrast, since F1 doesn’t include true negatives by definition, it would heavily penalize false predictions, which often aligns better with human evaluation of the performance. Therefore, even though F1 is not commonly used for the evaluation of image deconvolution tasks in natural images, we find that it has high utility for the data at hand in our application, especially if combined with additional metrics.

The main drawback of the F1 score is that it depends on a subjective binary threshold. In all experiments, binarization of the ground truth data was performed by a biologist. In cases in which publicly-available data was used for training (Fig. 2), we used the binarization performed by the original publication. CompbPlex predictions were used as outputted from the model with no further binarization.

#### Pearson correlation

Pearson correlation coefficients were computed by comparing the GT single-protein images with their recovered image using CombPlex for each FOV and each protein. For CODEX images, the images first underwent a 5×5 average filter to smooth the data. The objective was to perform patch-wise correlation between the reconstructed and the original image.

Pearson’s correlation coefficient (R) is widely used to evaluate image similarity in the medical imaging domain ^43–46^. It is highly advantageous in having no parameters. Unlike MSE or PSNR, it has a bounded range between −1 to 1, making it interpretable and intuitive. Moreover, we found that it was specifically adequate for the task at hand, because it is not sensitive to systematic linear changes between the ground truth and the reconstruction. Fluorescent channels have distinct properties, with some being inherently brighter than others ^47^. Similarly, different metal isotopes have different levels of sensitivity in mass-based imaging ^15^. As such, if one would measure the same protein using the same antibody, but once using channel X (e.g. ATTO-550) and once using channel Y (e.g. DYLight-751), the stainings will be correlated, but the values will not necessarily be the same, and systematic differences will be expected. For example, supplementary figure 10C shows the same antibody against pan-keratin, measured using two fluorophores: ATTO-550 and DYLight-751. A pixel-wise comparison between these two images shows that while the pixel values are correlated, they are not identical, and don’t fall around the Y=X line (**Supp. Fig. 10D**). Since in our evaluation of CombPlex the ground truth is not always measured using the same fluorescent channel or mass channel as the multiplexed image, we expect such systematic differences in the values between the ground truth and the image reconstructed by CombPlex. Therefore, Pearson correlation, which measures how well two images are linearly correlated, is particularly fitting for the properties of our data.

#### PSNR

PSNR was computed by comparing the GT single-protein images with their reconstructed image using CombPlex. This was done separately for each FOV and each protein using the following formula:

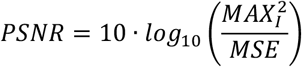

Where *MAX*_1_ is the theoretical maximal pixel value of the original image: 2^16^ for CODEX images and 2^8^ for MIBI-TOF images.

As mentioned above, PSNR is closely related to R and both are widely-accepted metrics for evaluation of image reconstruction tasks ^48^. Their major difference stems from the fact that the Pearson correlation estimates the linear correlation, while PSNR uses the maximal intensity of the signal to normalize the MSE. In our data, this adds another layer of complexity, because the dynamic range of fluorescent images is often lower than 2^16^, which is the theoretical maximal intensity for 16-bit images. Using 2^16^ as the maximum value for normalization may inflate the results, making it less meaningful. In our experience, this was also the case for sparse targets, making it hard to compare the reconstruction quality between targets.

#### SSIM

SSIM was computed by comparing the GT single-protein images with their reconstructed image using CombPlex for each FOV and each protein. It was calculated on 7×7 moving windows with the following formula:

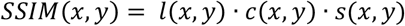

Where:

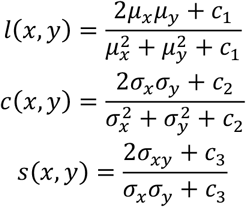

Where *c*_1_, *c*_2_ and *c*_3_ are positive constants used to avoid a null denominator.

SSIM is used to measure the similarity between two images. SSIM ranges between −1 to 1, where 1 indicates highly correlated images and −1 indicates anti correlated images, and higher is better. SSIM is considered to be correlated with the three quality attributes associated with the perception of the human eye: luminance, contrast and structure. Unlike PSNR and Pearson correlation, which rely on differences in expression values between the images, SSIM also accounts for expression variance differences. This allows SSIM to assign higher scores to images with a linear relationship, even if the values were not reconstructed exactly, a scenario that PSNR might penalize. Accounting for differences in variance is beneficial for the biological use case, where it is important to maintain the overall interrelations of image intensities during reconstruction rather than focusing solely on exact intensity values. On the other hand, in protein images with high sparsity, calculating with a sliding window can artificially inflate SSIM values, which might not accurately reflect the human perspective that focuses more on the actual signal in the image.

### Figure creation

Illustrations were created using Biorender templates (https://biorender.com).

## Data and code availability

The code for CombPlex can be found at: https://github.com/KerenLab/CombPlex. The data used in our experiments can be found at: https://www.ebi.ac.uk/biostudies/studies/S-BIAD873. List of reagents and instruments can be found in the table S1 under Key Resources.

## Acknowledgements

L.K. is supported by the Enoch, Azrieli, Sharon Levine and Rising Tide foundations, and grants funded by the Schwartz/Reisman Collaborative Science Program, the European Research Council (948811), the Israel Science Foundation (2481/20, 3830/21) within the Israel Precision Medicine Partnership program. I.M is supported by a EU - Horizon 2020 - MSCA Individual Fellowship (890733). S.B. is a Robin Chemers Neustein AI Fellow and acknowledges funds from the Carolito Stiftung and the NVIDIA Applied Research Accelerator Program. C.M.S is supported by the European Union (ERC, CAR-TIME, 101116768), the German Research Foundation (INST 37/1228-1 and INST 37/1302-1 FUGG, and Germany’s Excellence Strategy - EXC 2180 – 390900677), the Swiss Life Jubiläumsstiftung, the Mach-Gaensslen Stiftung Schweiz, the American Society of Hematology Research Restart Award, and the State of Baden-Württemberg within the Centers for Personalized Medicine Baden-Württemberg (ZPM). O.E. is supported by the Israel Cancer Research Foundation (ICRF). Phenocycler imaging was made possible thanks to the support of de Picciotto Cancer Cell Observatory In memory of Wolfgang and Ruth Lesser.

