## Supplementary figures for "Escalating High-dimensional Imaging using Combinatorial Channel Multiplexing and Deep Learning"

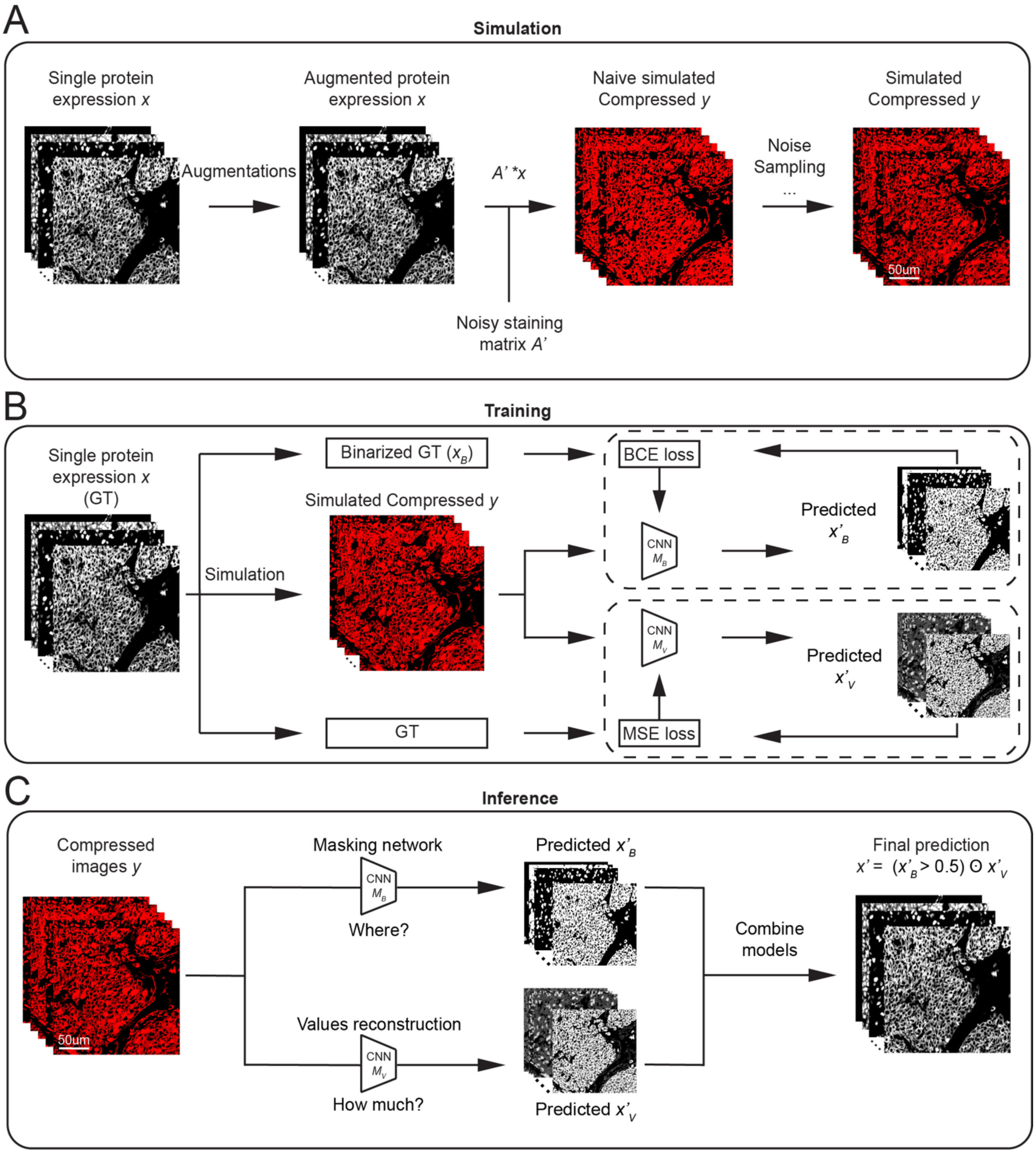


**Supplementary figure 1: Algorithmic overview of CombPlex**

**(A)** Illustration of the simulation process of compressed images. **(B)** Illustration of the training process involving simulated compressed images. BCE loss is calculated by comparing the binarized masks of the single-protein images and the binary mask predictions generated by the neural network, using the compressed images as input. MSE loss is calculated by comparing the single-protein images and the predictions generated by the neural network, using the compressed images as input. **(C)** Illustration of the inference procedure. Compressed images are provided as input to the masking and the value reconstruction networks. The prediction from both is intersected to produce the final prediction.

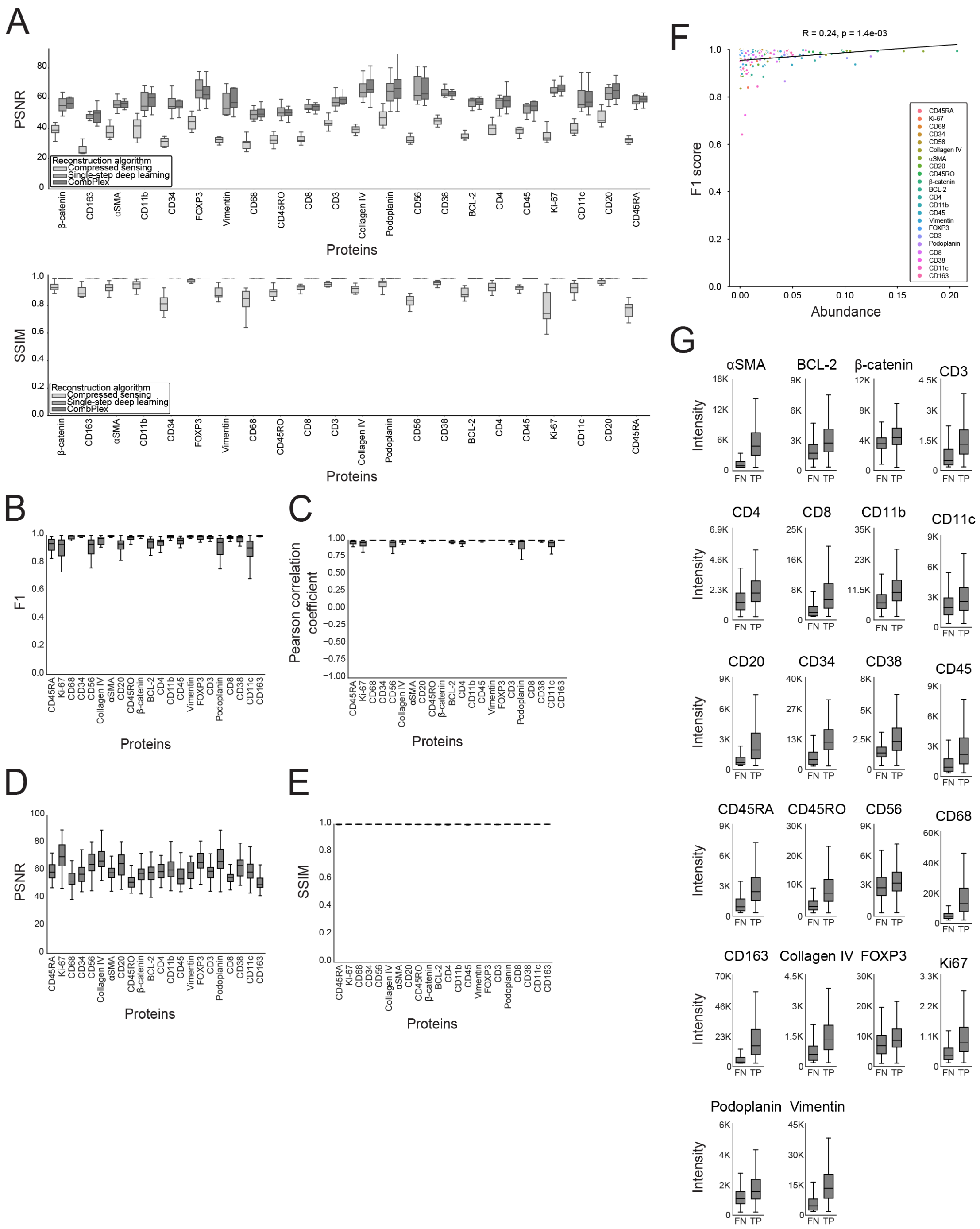


**Supplementary figure 2: Analysis of simulations**

The data from Schurch et al 1 was used to simulate combinatorically-compressed images. **(A)** PSNR and SSIM scores (y-axis) for each of the 22 proteins (x-axis) recovered by using compressed sensing, single-step deep learning or CombPlex. **(B-E)** Comparison between the ground truth images and the images recovered by CombPlex on the entire dataset using 4-fold cross validation. For each protein (x-axis) shown are the F1 scores (B), Pearson correlation (C), PSNR (D) and SSIM (E). **(F)** The correlation between the F1 score (y-axis) of different proteins and their abundance (x-axis) on a test set of 8 FOVs. **(G)** Intensity values in ground truth images (y-axis) of FN and TP pixels (x-axis) for 22 proteins on a test set of 8 FOVs. FN errors tend to have lower intensities than TP pixels.

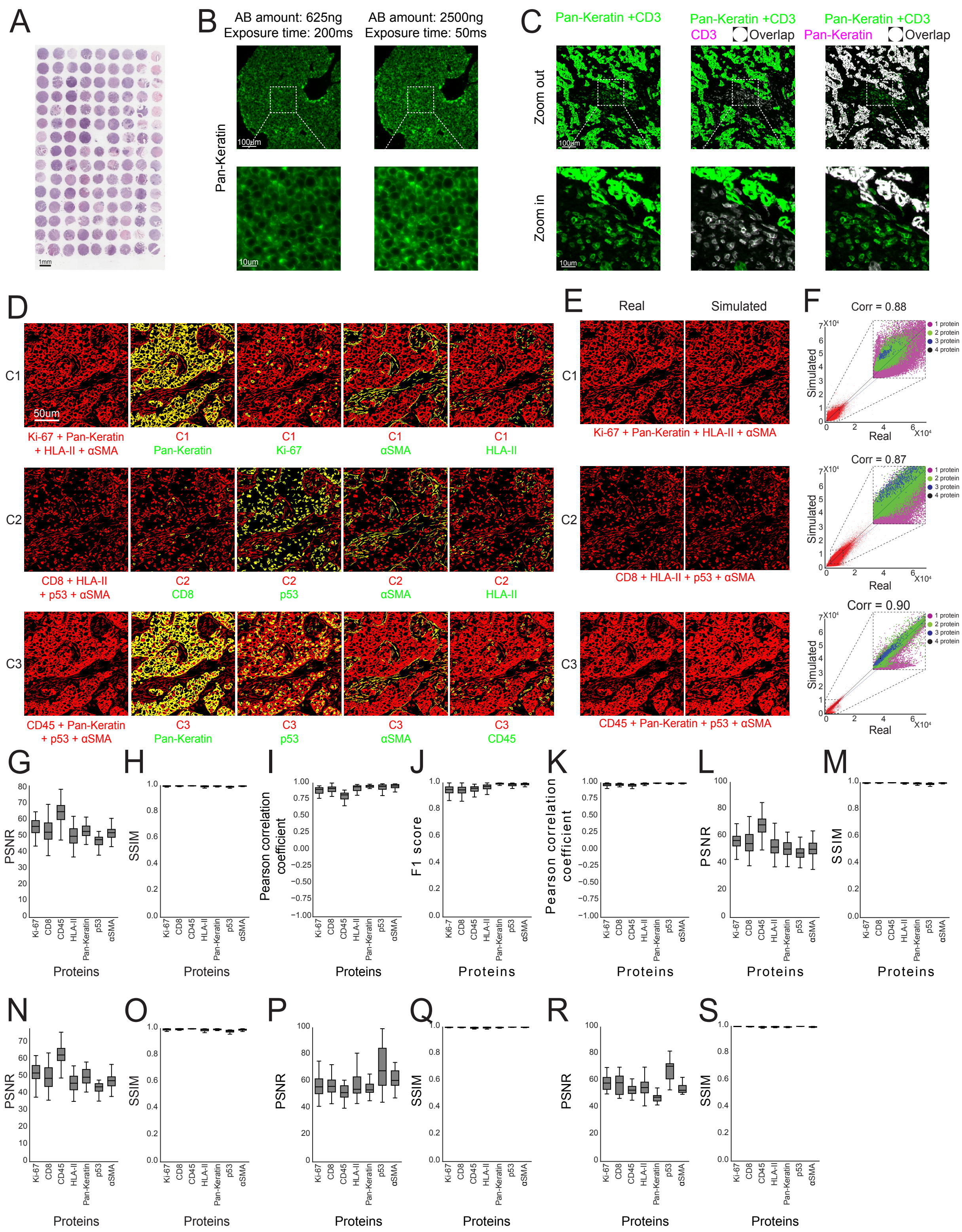

A
B
AB amount: 625ng
Exposure time: 200ms
AB amount: 2500ng
Exposure time: 50ms
Pan-Keratin
C
Pan-Keratin +CD3
Pan-Keratin +CD3
CD3
Overlap
Pan-Keratin +CD3
Pan-Keratin
Overlap
Zoom out
Zoom in
D
Ki-67 + Pan-Keratin
+ HLA-II + αSMA
C1 Pan-Keratin
C1 Ki-67
C1 αSMA
C1 HLA-II
CD8 + HLA-II
+ p53 + αSMA
C2 CD8
C2 p53
C2 αSMA
C2 HLA-II
CD45 + Pan-Keratin
+ p53 + αSMA
C3 Pan-Keratin
C3 p53
C3 αSMA
C3 CD45
E
Real
Simulated
Ki-67 + Pan-Keratin + HLA-II + αSMA
CD8 + HLA-II + p53 + αSMA
CD45 + Pan-Keratin + p53 + αSMA
F
Corr = 0.88
Corr = 0.87
Corr = 0.90
1 protein
2 protein
3 protein
4 protein
Simulated
Real
G
H
I
J
K
L
M
N
O
P
Q
R
S
PSNR
SSIM
Pearson correlation coefficient
F1 score
Proteins
Ki-67
CD8
CD45
HLA-II
Pan-Keratin
p53
αSMA

**Supplementary figure 3: Evaluation of CombPlex on small-scale CODEX experiments**
**(A)** A TMA of 132 breast carcinoma cores. Samples were used to generate compressed and ground truth images. **(B)** Example images of Pan-Keratin stained using different concentrations and exposure times. Antibodies on the same channel are tittered to accommodate the same exposure without loss of signal. **(C)** An example image of Pan-Keratin and CD3 measured together on the same channel (green) or separately (magenta). White pixels indicate overlap. **(D)** Overlay of seven experimentally-measured single-protein images (ground truth, green) with the respective compressed images that contain all of their signals (red). Yellow pixels indicate overlap. **(E)** Side-by-side comparison of experimentally compressed images (left) and simulated compressed images (right). **(F)** Shown is the correlation between the experimentally-compressed and simulated compressed images shown in (C). Inset is colored by the number of overlapping proteins per pixel. **(G)** PSNR scores (y-axis) for each protein (x-axis) recovered by CombPlex on 32 test FOVs. The models were trained on experimentally compressed images. **(H)** Same as (G), showing the SSIM scores between the reconstructed and ground truth images. **(I)** Pearson correlation coefficients (y-axis) for each protein (x-axis) calculated on the intersection of pixels between the reconstructed and ground truth images. **(J)** F1 scores (y-axis) for each protein (x-axis) recovered by CombPlex using 4-fold cross validation. **(K)** Same as (J), showing the Pearson correlation coefficient between the reconstructed and ground truth images. **(L)** Same as (J), showing the PSNR between the reconstructed and ground truth images. **(M)** Same as (J), showing the SSIM between the reconstructed and ground truth images. **(N)** PSNR scores (y-axis) for each protein (x-axis) recovered by CombPlex on 32 test FOVs. The models were trained on simulated compressed images. **(O)** Same as (N), showing the SSIM scores between the reconstructed and ground truth images. **(P)** CombPlex was trained on experimentally compressed images of breast, lung and colon carcinoma. Shown are the PSNR scores (y-axis) for each protein (x-axis) recovered by CombPlex on 44 breast, lung and colon carcinoma test FOVs. **(Q)** Same as (P), showing the SSIM between the reconstructed and ground truth images. **(R-S)** Same as (P-Q) when the model trained on breast, lung and colon carcinoma was evaluated on 20 PDAC samples.

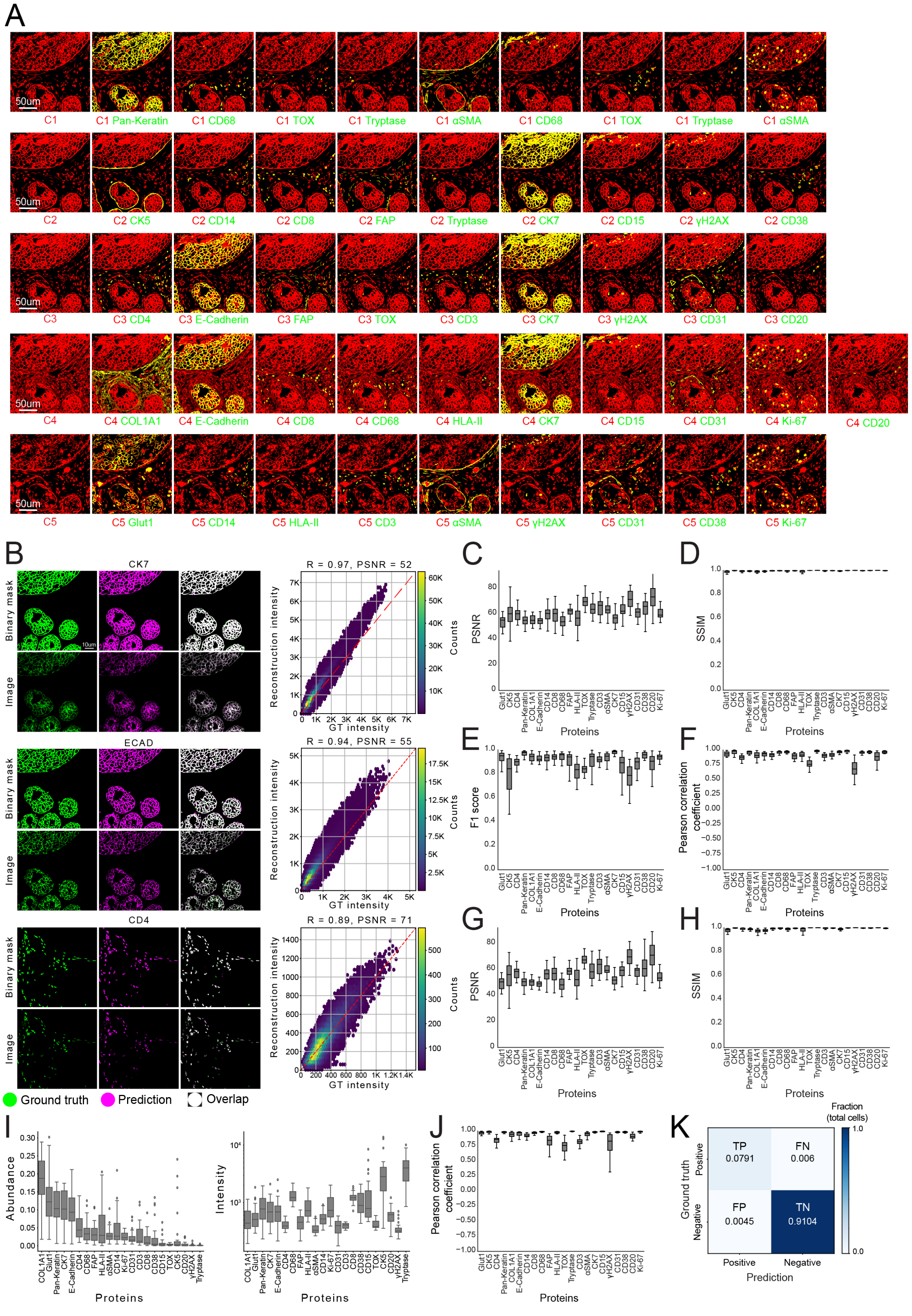

A
50um
C1
C1 Pan-Keratin
C1 CD68
C1 TOX
C1 Tryptase
C1 αSMA
C1 CD68
C1 TOX
C1 Tryptase
C1 αSMA
C2
C2 CK5
C2 CD14
C2 CD8
C2 FAP
C2 Tryptase
C2 CK7
C2 CD15
C2 γH2AX
C2 CD38
C3
C3 CD4
C3 E-Cadherin
C3 FAP
C3 TOX
C3 CD3
C3 CK7
C3 γH2AX
C3 CD31
C3 CD20
C4
C4 COL1A1
C4 E-Cadherin
C4 CD8
C4 CD68
C4 HLA-II
C4 CK7
C4 CD15
C4 CD31
C4 Ki-67
C4 CD20
C5
C5 Glut1
C5 CD14
C5 HLA-II
C5 CD3
C5 αSMA
C5 γH2AX
C5 CD31
C5 CD38
C5 Ki-67
B
CK7
ECAD
CD4
Binary mask
Image
10um
R = 0.97, PSNR = 52
R = 0.94, PSNR = 55
R = 0.89, PSNR = 71
Reconstruction intensity
GT intensity
Counts
Ground truth
Prediction
Overlap
C
D
E
F
G
H
PSNR
SSIM
F1 score
Pearson correlation coefficient
Proteins
Glut1
CK5
CD4
Pan-Keratin
COL1A1
E-Cadherin
CD14
CD8
CD68
FAP
HLA-II
TOX
Tryptase
CD3
αSMA
CK7
CD15
γH2AX
CD31
CD38
CD20
Ki-67
I
Abundance
Intensity
J
K
Fraction (total cells)
TP
0.0791
FN
0.006
FP
0.0045
TN
0.9104
Ground truth
Positive
Negative
Prediction

**Supplementary figure 4: Evaluation of CombPlex on large-scale CODEX experiments**

**(A)** Overlay of 22 experimentally-measured single-protein images (ground truth, green) with the respective compressed images that contain all their signals (red). Yellow pixels indicate overlap. **(B)** Left: Shown are images of ground truth (green), CombPlex prediction (magenta) and their overlay (white) of CK7, ECAD and CD4 (left). For each protein, the top row shows the binary mask, and the bottom row shows the values. Right: Pixel-wise correlation between the ground truth and prediction intensities (right). **(C-D)** CombPlex was trained on experimentally compressed images. For each protein (x-axis) shown are the PSNR (C) or SSIM (D) between the reconstructed and ground truth images. **(E-H)** CombPlex was trained on compressed images generated by simulations in silico and evaluated on 30 test FOVs. For each protein (x-axis) shown are the F1 scores (E) Pearson correlation (F) PSNR (G) or SSIM (H) between the 30 reconstructed and ground truth images. **(I)** Left: abundance (y-axis), calculated as the number of positive pixels, for each protein (x-axis). Right: median intensity values (y-axis) for each protein (x-axis). **(J)** Pearson correlation coefficients (y-axis) for each protein (x-axis) calculated only on the intersection of pixels between the reconstructed and ground truth images. **(K)** For each protein in the 30 test FOVs, each pixel was compared between the ground truth images and the images reconstructed by CombPlex. Pixels were classified as TP, FN, FP and TN.

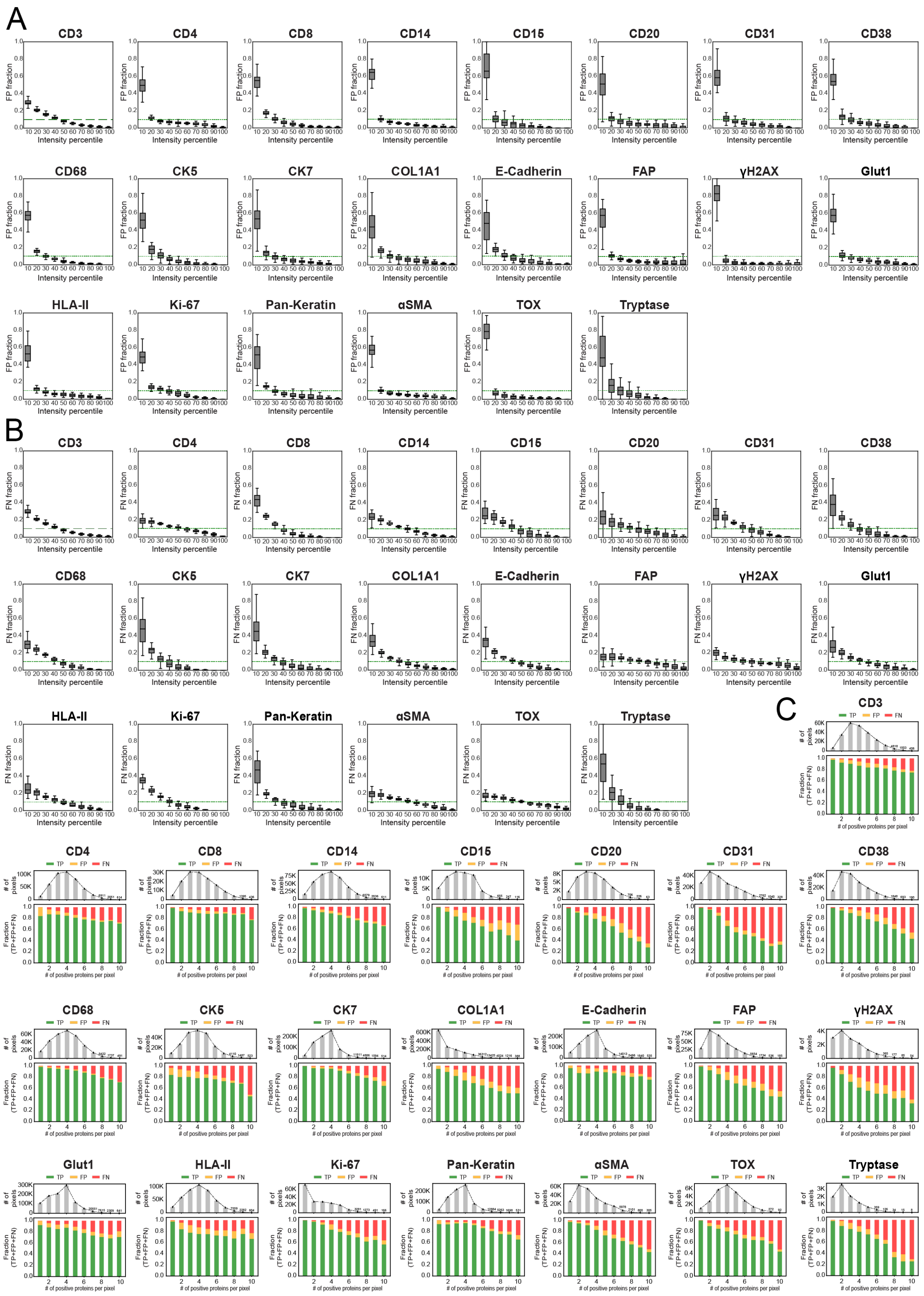
A
CD3
CD4
CD8
CD14
CD15
CD20
CD31
CD38
CD68
CK5
CK7
COL1A1
E-Cadherin
FAP
γH2AX
Glut1
HLA-II
Ki-67
Pan-Keratin
αSMA
TOX
Tryptase
FP fraction
Intensity percentile
B
FN fraction
C
TP
FP
FN
### of pixels
Fraction (TP+FP+FN)
### of positive proteins per pixel

**Supplementary figure 5: False Negative and False Positive analysis in large-scale CODEX CombPlex experiments**
CODEX cyclic imaging was used to image 22 proteins on a single tissue section, both compressed, and individually. For each protein are shown: **(A)** Shown are the fractions of FP pixels (y-axis) for each decile of GT intensities (x-axis) for all distinct single-protein images across all FOVs. The green line indicates the 0.1 expectation from a uniform distribution. **(B)** Shown are the fractions of FN pixels (y-axis) for each decile of GT intensities (x-axis) for all distinct single-protein images across all FOVs. The green line indicates the 0.1 expectation from a uniform distribution. **(C)** Bottom: Shown is the composition of TP (True Positives), FP (False Positives), and FN (False Negatives) pixels (y-axis) across all distinct single-protein images of all FOVs as a function of the number of proteins that are positive in each pixel (x-axis). Top: The number of pixels corresponding to each protein count.

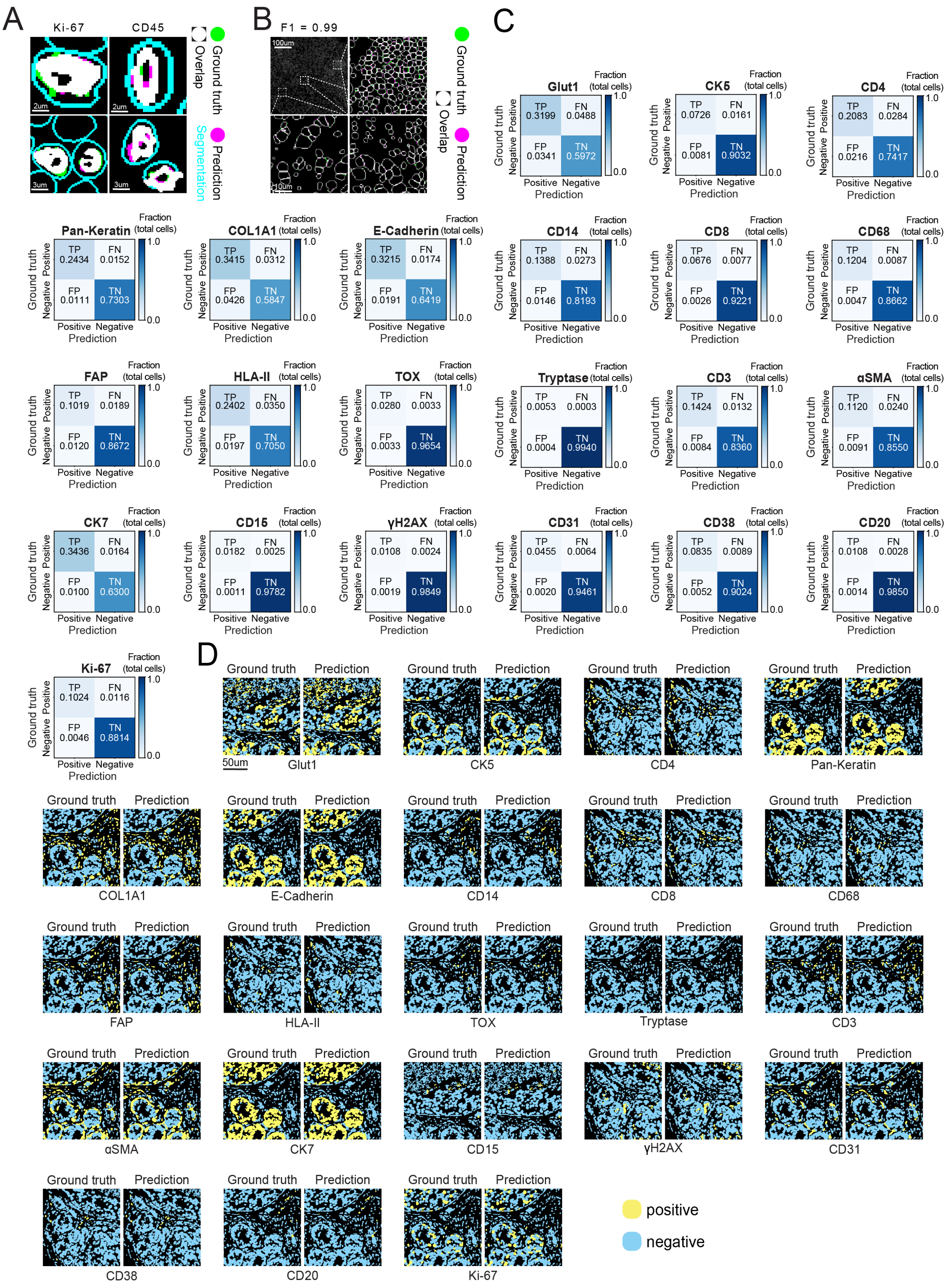

A
Ki-67
CD45
Ground truth
Overlap
Prediction
Segmentation
2um
2um
3um
3um
B
F1 = 0.99
100um
10um
Ground truth
Overlap
Prediction
C
Glut1 Fraction (total cells)
TP 0.3199 FN 0.0488
FP 0.0341 TN 0.5972
CK5 Fraction (total cells)
TP 0.0726 FN 0.0161
FP 0.0081 TN 0.9032
CD4 Fraction (total cells)
TP 0.2083 FN 0.0284
FP 0.0216 TN 0.7417
Pan-Keratin Fraction (total cells)
TP 0.2434 FN 0.0152
FP 0.0111 TN 0.7303
COL1A1 Fraction (total cells)
TP 0.3415 FN 0.0312
FP 0.0426 TN 0.5847
E-Cadherin Fraction (total cells)
TP 0.3215 FN 0.0174
FP 0.0191 TN 0.6419
CD14 Fraction (total cells)
TP 0.1388 FN 0.0273
FP 0.0146 TN 0.8193
CD8 Fraction (total cells)
TP 0.0676 FN 0.0077
FP 0.0026 TN 0.9221
CD68 Fraction (total cells)
TP 0.1204 FN 0.0087
FP 0.0047 TN 0.8662
FAP Fraction (total cells)
TP 0.1019 FN 0.0189
FP 0.0120 TN 0.8672
HLA-II Fraction (total cells)
TP 0.2402 FN 0.0350
FP 0.0197 TN 0.7050
TOX Fraction (total cells)
TP 0.0280 FN 0.0033
FP 0.0033 TN 0.9654
Tryptase Fraction (total cells)
TP 0.0053 FN 0.0003
FP 0.0004 TN 0.9940
CD3 Fraction (total cells)
TP 0.1424 FN 0.0132
FP 0.0084 TN 0.8360
αSMA Fraction (total cells)
TP 0.1120 FN 0.0240
FP 0.0091 TN 0.8550
CK7 Fraction (total cells)
TP 0.3436 FN 0.0164
FP 0.0100 TN 0.6300
CD15 Fraction (total cells)
TP 0.0182 FN 0.0025
FP 0.0011 TN 0.9782
γH2AX Fraction (total cells)
TP 0.0108 FN 0.0024
FP 0.0019 TN 0.9849
CD31 Fraction (total cells)
TP 0.0455 FN 0.0064
FP 0.0020 TN 0.9461
CD38 Fraction (total cells)
TP 0.0835 FN 0.0089
FP 0.0052 TN 0.9024
CD20 Fraction (total cells)
TP 0.0108 FN 0.0028
FP 0.0014 TN 0.9850
Ki-67 Fraction (total cells)
TP 0.1024 FN 0.0116
FP 0.0046 TN 0.8814
Ground truth Positive Negative
Prediction Positive Negative
1.0
0.0
D
Ground truth
Prediction
50um
Glut1
CK5
CD4
Pan-Keratin
COL1A1
E-Cadherin
CD14
CD8
CD68
FAP
HLA-II
TOX
Tryptase
CD3
αSMA
CK7
CD15
γH2AX
CD31
CD38
CD20
Ki-67
positive
negative

**Supplementary figure 6: Classification analysis in large-scale CODEX CombPlex experiments**

**(A)** Shown are example zoomed-in images of ground truth (green) and prediction (magenta) of Ki-67 (left) and CD45 (right). White pixels indicate overlap. Segmentation is shown in cyan. **(B)** Segmentation masks generated by DeepCell using DAPI and ground truth (green) or prediction (magenta) membrane channels. White pixels indicate overlap. (C-D) CODEX cyclic imaging was used to image 22 proteins on a single tissue section, both combinatorically-compressed, and individually. For each protein (rows), cells were classified as positive or negative based on either the ground truth images, or the images reconstructed by CombPlex. **(C)** Shown is the confusion between both classification schemes across 30 test FOVs. TP: True Positive, FN: False Negative, FP: False Positive and TN: True Negative. **(D)** A representative image showing cells classified as positive (yellow) and negative (cyan) for each protein (rows) using ground truth (left) or reconstructed (right) images.

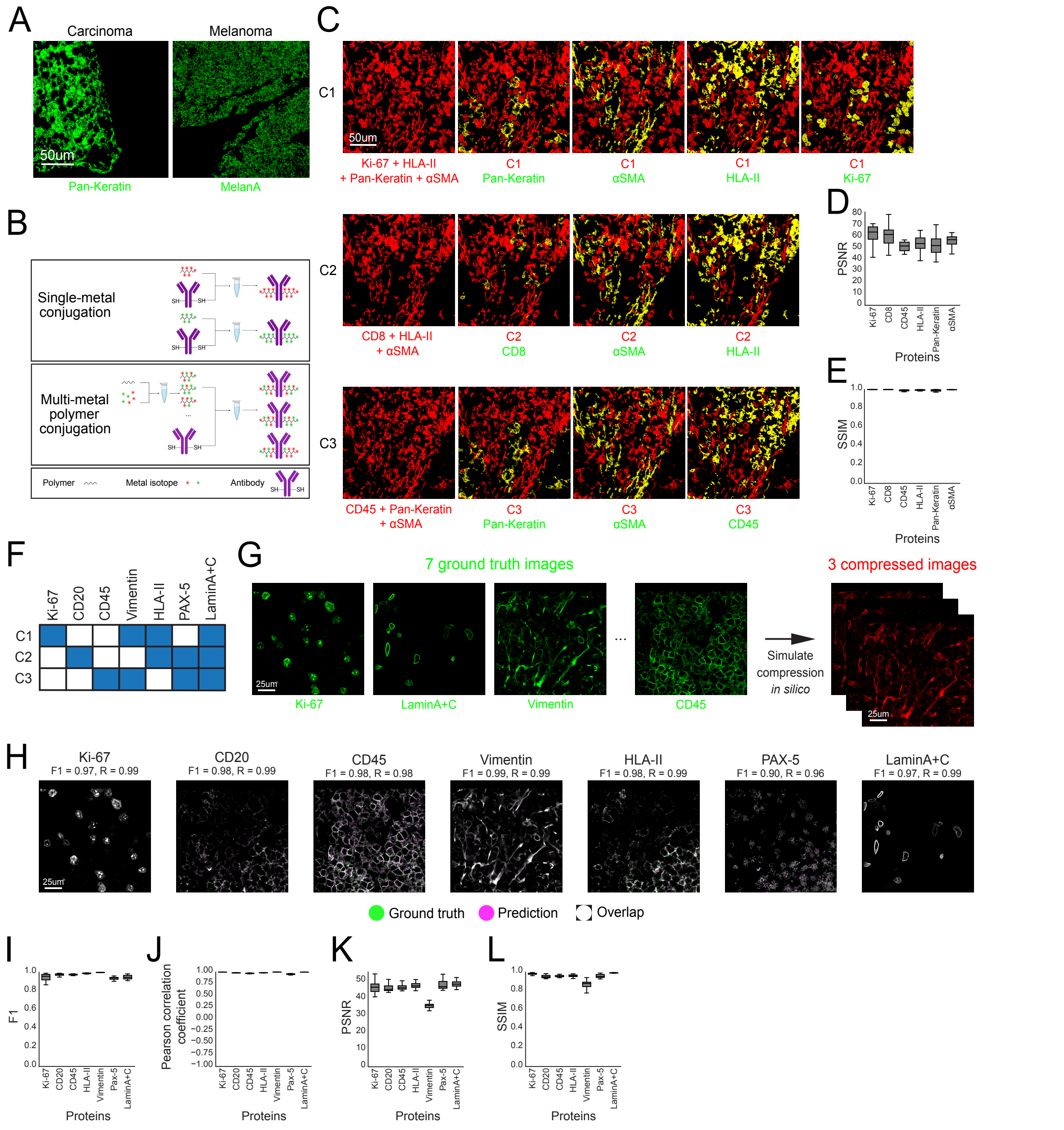


**Supplementary figure 7: Evaluation of MIBI-TOF CombPlex experiments**

**(A)** Shown are example images of Pan-Keratin and MelanA signals in breast carcinoma and melanoma metastases respectively. **(B)** Illustration of metal-conjugation methods. Single-metal conjugation (top): separate lots of the same antibody are reduced and conjugated separately, each to a single-metal labeled polymer. Multi-metal polymer conjugation (bottom): Unlabeled polymers are loaded with multiple isotopes and conjugated to a reduced antibody. **(C)** Overlay of six experimentally-measured single-protein images (ground truth, green) with the respective compressed images that contain all of their signals (red). Yellow pixels indicate overlap. **(D)** PSNR scores (y-axis) for each protein (x-axis) recovered by CombPlex on 19 test FOVs. The model was trained on compressed images generated by simulations on previously-available data. **(E)** Same as (D), showing SSIM scores between the reconstructed and ground truth images. **(F)** The compression matrix employed to condense the data from Bai et al.

**(G)** *In silico* compression of 7 protein images into 3 combinatorically-compressed images. **(H)** CombPlex reconstruction (magenta) is overlaid on the ground truth single-protein image (green). White pixels indicate overlap. **(I-L)** For each protein (x-axis) shown are the F1 scores (I) Pearson correlation (J) PSNR (K) or SSIM (L) between the 18 reconstructed and ground truth images.

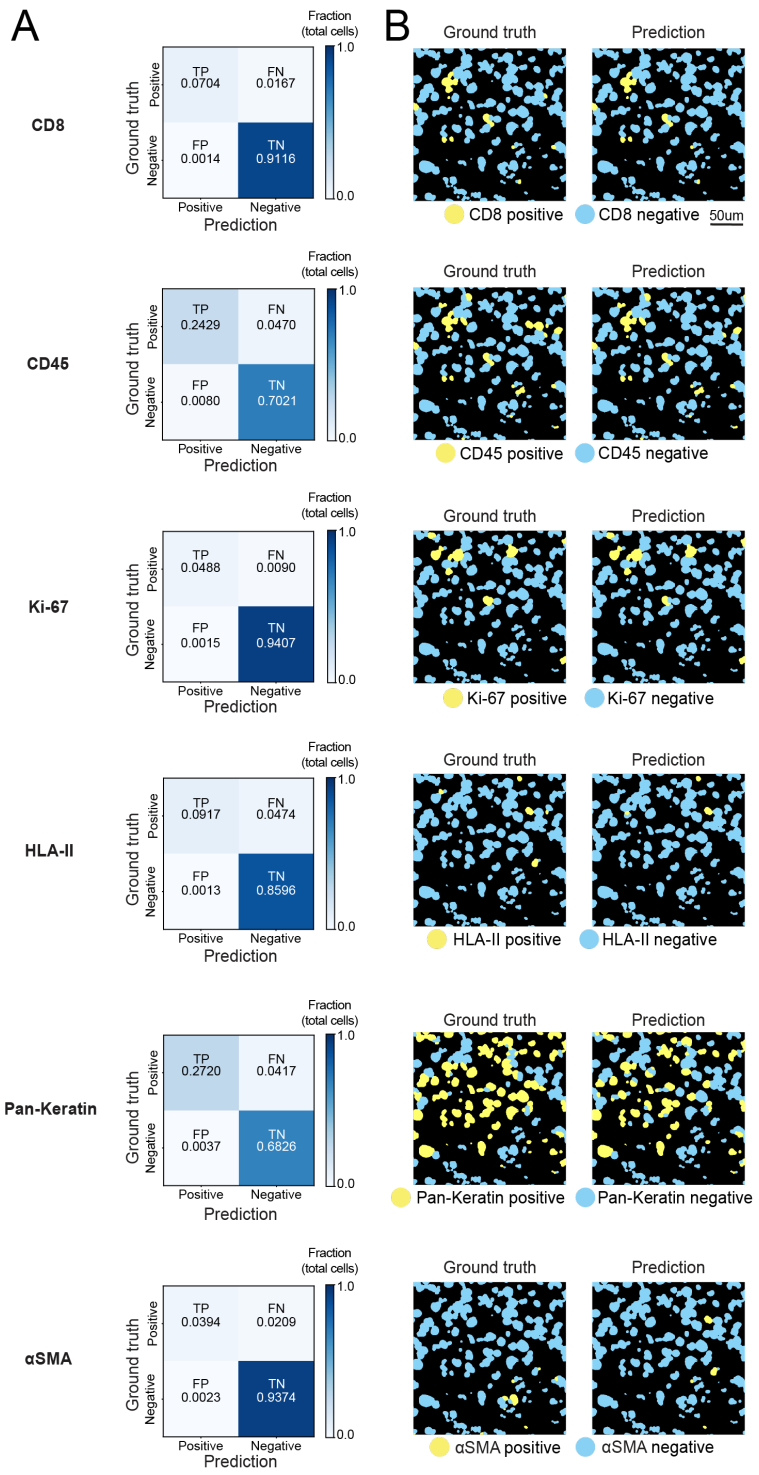


**Supplementary figure 8: Classification analysis in MIBI-TOF CombPlex experiments** MIBI-TOF was used to image six proteins on a single tissue section, both combinatorically-compressed, and individually. For each protein (rows), cells were classified as positive or negative based on either the ground truth or reconstructed single-protein image. **(A)** Shown is the confusion between both classification schemes across 19 FOVs. TP: True Positive, FN: False Negative, FP: False Positive and TN: True Negative. **(B)** Cells classified as positive (yellow) and negative (cyan) for each protein (rows) using ground truth (left) or reconstructed (right) images.

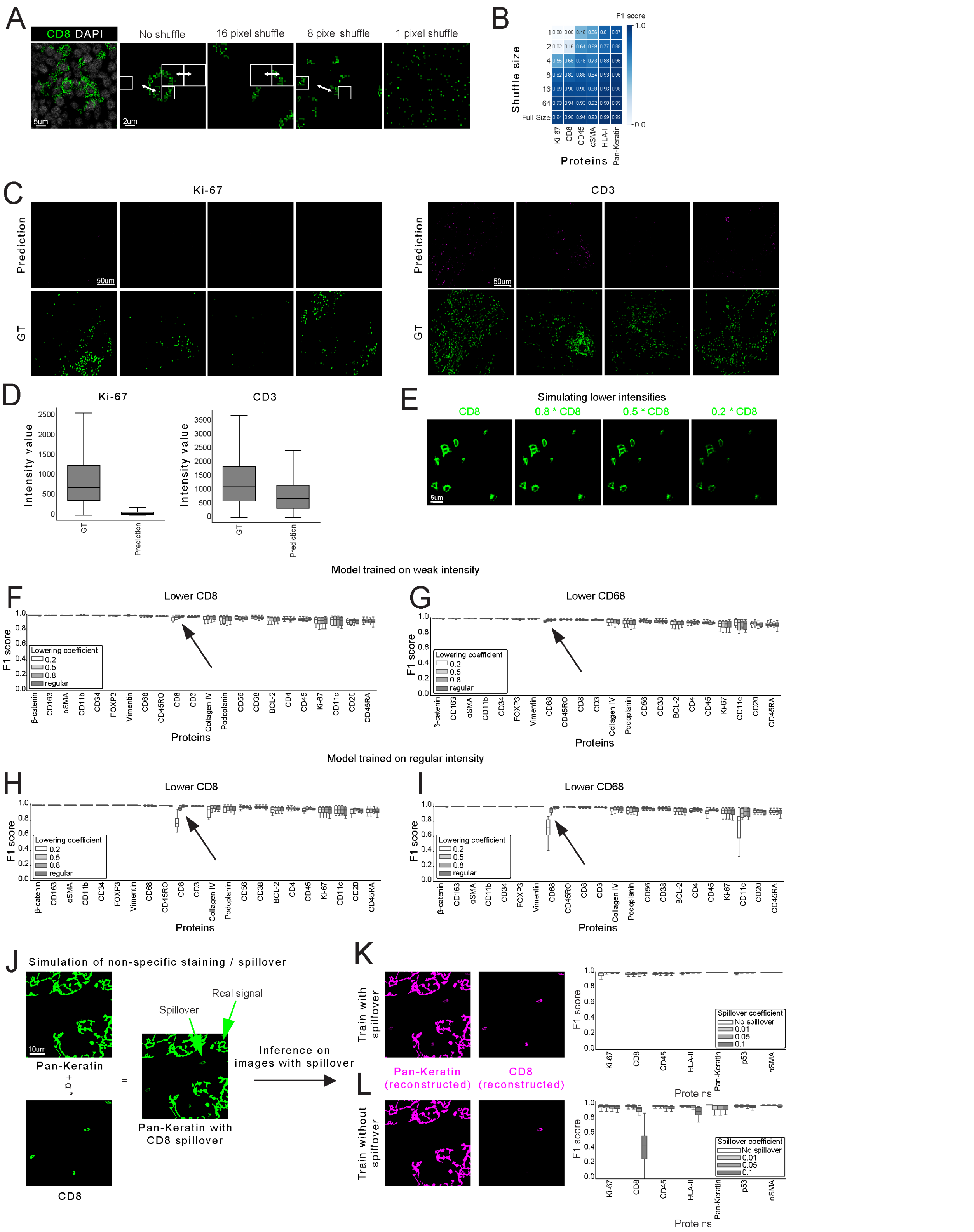


**Supplementary figure 9: Exploration of the performance of CombPlex**

**(A)** An example MIBI-TOF image of CD8 (green) and dsDNA (white). Images were shuffled using patches of varying sizes. **(B)** CombPlex was trained on regular MIBI-TOF data and then evaluated on data shuffled with different patch sizes. Shown are the median F1 scores for each protein (x-axis) on the data shuffled with different patch sizes (y-axis). **(C)** Left: Example images of the prediction (top) and ground truth (bottom) images for Ki-67, in simulations in which it appeared in the training, but was dropped out from the test set. Right: Same for CD3. **(D)** Left: Intensity values (y-axis) of ground truth Ki-67 images, and images predicted by CombPlex for Ki-67 when it is dropped out of the experiment. Right: Same for CD3. **(E)** An example image of CD8 at original intensity, 80% intensity, 50% intensity and 20% intensity. **(F)** F1 scores (y-axis) for all proteins (x-axis) recovered by CombPlex evaluated on images with different intensities of CD8 (colored bars). CombPlex was trained on images with corresponding weak CD8 intensity. **(G)** Same as (F) for CD68. **(H)** Same as (F), but CombPlex was trained on original intensity images. **(I)** Same as (G), but CombPlex was trained on original intensity images. **(J)** Non-specific staining of Pan-Keratin was simulated by adding to it the CD8 signal in varying intensities. **(K)** Left: Examples of CombPlex predictions for Pan-Keratin and CD8 when training with spillover. Right: F1 scores (y-axis) for all proteins (x-axis) recovered by CombPlex. **(L)** Same as K, but when training CombPlex on images with no spillover.

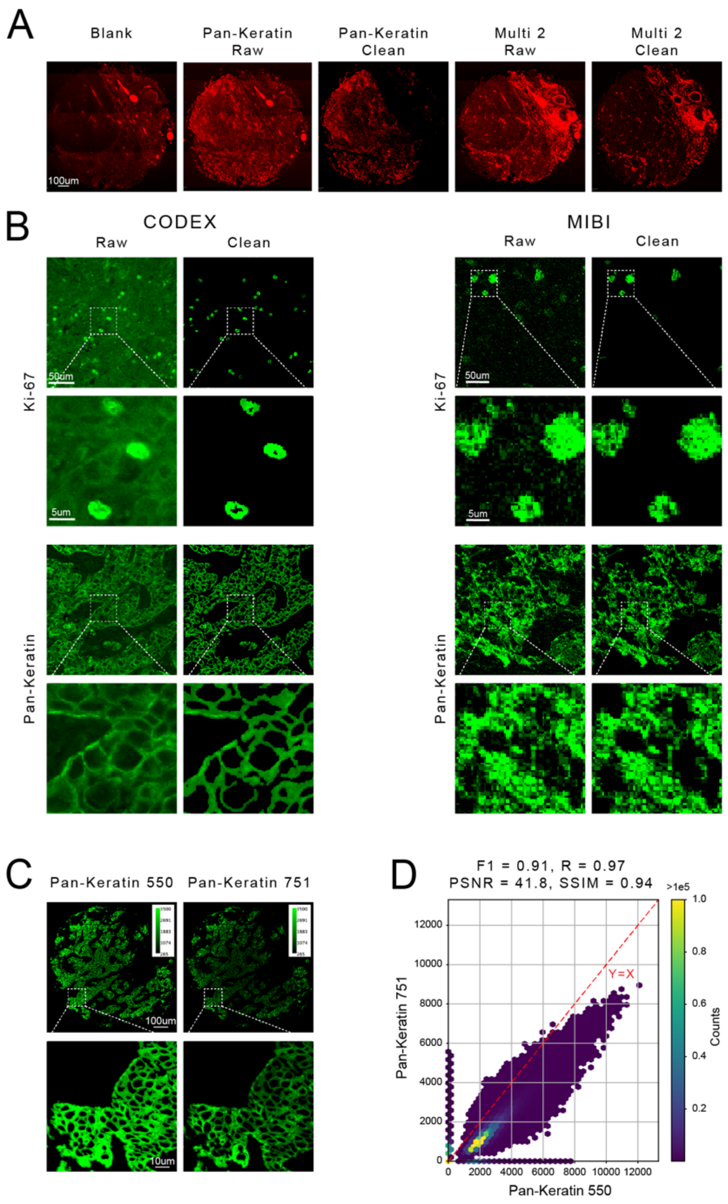


**Supplementary figure 10: Methods**

**(A)** Removal of autofluorescence as done automatically by the Phenocycler Fusion instrument. Shown is an example image of a blank cycle (left), used to remove autofluorescence from raw single (middle) or compressed (right) images. **(B)** Shown are example images of Ki-67 (Top) and Pan-Keratin (Bottom) before and after cleaning by using a pixel classifier on CODEX (left) and MIBI (right) images. **(C)** Shown are example images of Pan-Keratin measured using either ATTO550 (left) or DYLight 751 (right). **(D)** Shown is the correlation between Pan-Keratin measure using ATTO550 and DYLight751.
